# Integrative multi-omic sequencing reveals the MMTV-Myc mouse model mimics human breast cancer heterogeneity

**DOI:** 10.1101/2023.03.28.534611

**Authors:** Carson D. Broeker, Mylena M. O. Ortiz, Michael S. Murillo, Eran R. Andrechek

## Abstract

**Background:** Breast cancer is a complex and heterogeneous disease with distinct subtypes and molecular profiles corresponding to different clinical outcomes. Mouse models of breast cancer are widely used, but their relevance in capturing the heterogeneity of human disease is unclear. Previous studies have shown the heterogeneity at the gene expression level for the MMTV-Myc model, but have only speculated on the underlying genetics.

**Results:** Herein, we examine three common histological subtypes of the MMTV-Myc model through whole genome sequencing and have integrated these results with gene expression data. Significantly, key genomic alterations driving cell signaling pathways were well conserved within histological subtypes. Genomic changes included frequent, co-occurring mutations in KIT and RARA in the microacinar histological subtype as well as SCRIB mutations in the EMT subtype. EMT tumors additionally displayed strong KRAS activation signatures downstream of genetic activating events primarily ascribed to KRAS activating mutations, but also FGFR2 amplification. Analogous genetic events in human breast cancer showed stark decreases in overall survival. In further analyzing transcriptional heterogeneity of the MMTV-Myc model, we report a supervised machine learning model that classifies MMTV-Myc histological subtypes and other mouse models as being representative of different human intrinsic breast cancer subtypes.

**Conclusions:** We conclude the well-established MMTV-Myc mouse model presents further opportunities for investigation of human breast cancer heterogeneity.

## Background

MYC is a master transcriptional regulator that is amplified in approximately 15–20% of breast cancers and overexpressed in up to 35% of breast cancers^1^. All MYC family genes (c-MYC, N-MYC, L-MYC, B-MYC) contain basic helix-loop-helix (bHLH) domains which allow for heterodimerization with MYC Associated Factor X (MAX) to bind consensus E-box sequence motifs on core gene promoter regions^2^; this allows for the initiation of transcription of MYC responsive genes, including regulation of proliferation, cell growth, differentiation, nucleotide biosynthesis, DNA replication, RNA levels, and apoptosis^3,4^. MYC amplification is often enriched in high grade breast cancers^5,6^, triple-negative breast cancers^7^, and basal-like breast cancers^8^. This is supported by evidence showing that MYC occupies the promoters of most active genes in tumorigenesis and that MYC acts as a non-specific amplifier of whole cell gene expression^9^. Triple-negative breast cancer (TNBC) and basal-like breast cancer subtypes carry poor prognostic clinical outcomes relative to other breast cancer subtypes^10^. In mouse models, several groups have demonstrated that overexpression of MYC alone is sufficient for the development of spontaneous mammary tumors with high penetrance^11–13^.

There is a clear, concerted interest to develop mouse models of breast cancer that recapitulate the heterogeneous features of human breast cancer to better understand disease dynamics. In response, numerous mouse models of breast cancer have been created, characterized by a wide variety of genetic drivers including constitutive overexpression of endogenous oncogenes specifically in the mouse mammary gland (MMTV-Neu, MMTV-Cyclin D1, MMTV-Akt1^14^), conditional overexpression of oncogenes in the mammary gland (MMTV-rtTA/TetO-NeuNT^15^, MTB/TWNT^16^), nonfunctional or conditional loss of tumor suppressor genes (Stat1^-/-^, MMTV-Cre/BRCA1^fl/fl^)^14^, or overexpression of exogenous transforming oncoproteins (MMTV-PyMT^17^). Various mouse models of breast cancer have had their gene expression profiles characterized^18,19^, with most models clustering largely within one human intrinsic breast cancer subtype. While many mouse models have been generated that model a specific aspect of human breast cancer biology, few realize the full spectrum of heterogeneity present within human breast tumors or accurately model clinical features^20^. Previous studies revealed the MMTV-Myc model of breast cancer mimics many human disease parameters, including substantial histological and transcriptional heterogeneity^13^. These analyses revealed a close association between the transcriptional profile and histological subtype among MMTV-Myc tumors. Other studies have also shown that tumors derived from different mouse models that share the same histological patterns cluster more tightly transcriptionally than they do with tumors of the same genotype with different histological features^18^. Importantly, MMTV-Myc samples have one of the most varied gene expression patterns amongst models, with distinct subsets of MMTV-Myc samples clustering closely with all intrinsic human breast cancer subtypes^18^.

While the contributions of the tumor microenvironment, epigenetics, tumor metabolome, immune composition, and other factors have been highlighted recently for their roles in cancer^21–23^, most often cancers are driven by genetic aberrations by activation of oncogenes and inactivation of tumor suppressor genes. The advent of anti-estrogenic compounds such as tamoxifen in the treatment of hormone-receptor positive breast cancers^24^ and the monoclonal antibody trastuzumab to treat HER2+ breast cancer^25^ have demonstrated the utility in targeting breast cancer with therapies based on genomic, transcriptomic, and immunohistochemical data. In realization of the complex molecular aberrations driving all cancers, a concerted effort in the form of The Cancer Genome Atlas (TCGA) was created and serves as a critical repository of integrated molecular cancer data for researchers^26^. Despite the clear relevance of genetic aberrations in human breast cancer progression and the wide use of mouse models to study human breast cancer *in vivo*, relatively few mouse models of breast cancer have been characterized at the genome level, barring MMTV-PyMT^27,28^, MMTV-Neu^27^, and NRL-PRL^29^ models. Previous mouse model whole genome sequencing (WGS) has revealed important human disease parallels, with sequencing of the MMTV-Neu mouse model revealing a conserved coamplification event that exists in 25% of human HER2+ breast cancers and 9% of all breast cancer. Preclinical functional studies showed the presence of this coamplification event increases migration *in vitro*, increases metastasis *in vivo*, and reduces distant metastasis free survival in humans^27^. In the MMTV-PyMT model, it was found that mutations in the phosphatase PTPRH led to increased phosphorylated EGFR levels and increased EGFR signaling^27^, with CRISPR ablation and rescue experiments demonstrating that PTPRH was normally dephosphorylating EGFR^30^. In each model, genomic alterations drove key aspects of cellular signaling, altering tumor biology, with key similarities to human cancer. These studies underscore the importance of characterizing genomic alterations in mouse models of cancer and the utility of integrating gene expression and DNA sequencing in GEMMs to find clinical parallels in human cancers.

Here, we report our findings of interrogating the heterogeneity within the MMTV-Myc mouse model. We performed WGS on three common, distinct histological subtypes that arose in this model. We hypothesized genomic heterogeneity would be present across subtypes but would be conserved within each histological subtype. Importantly, each of the subtypes have numerous tumors that have been previously analyzed for gene expression^13^. Integration of both genomic and transcriptomic data enabled the identification of active signaling pathways exploited in each histological subtype and their respective genetic origins. From this, targeted *in vivo* preclinical studies can be designed for the highest potential clinical translation into humans.

## Results

### Genomic analyses reveal conserved copy number gains in microacinar tumors

Based on the diverse transcriptional and histological subtypes observed in the MMTV-Myc tumors^13^, we hypothesized these phenotypes were due to a divergence in genomic changes conserved between each histological subtype. To ascertain putative conserved genomic changes within each histological subtype, short read WGS was performed on randomly selected tumors of microacinar, squamous, and EMT histological subtypes, later integrated with gene expression data obtained from matched samples as well as additional tumors from each subtype (**Figure 1**). A great deal of genomic heterogeneity was observed between the histological subtypes as shown by representative Circos plots for the microacinar (**Figure 2A**), squamous (**Figure 2B**), and EMT (**Figure 2C**) tumors. Few inversions and translocations were called across all tumors, consistent with rates of large structural rearrangements in human breast cancer^31^. Most differences in genetic aberrations between subtypes were confined to single nucleotide variants (SNVs) and copy number variants (CNVs).

**Figure 1:**
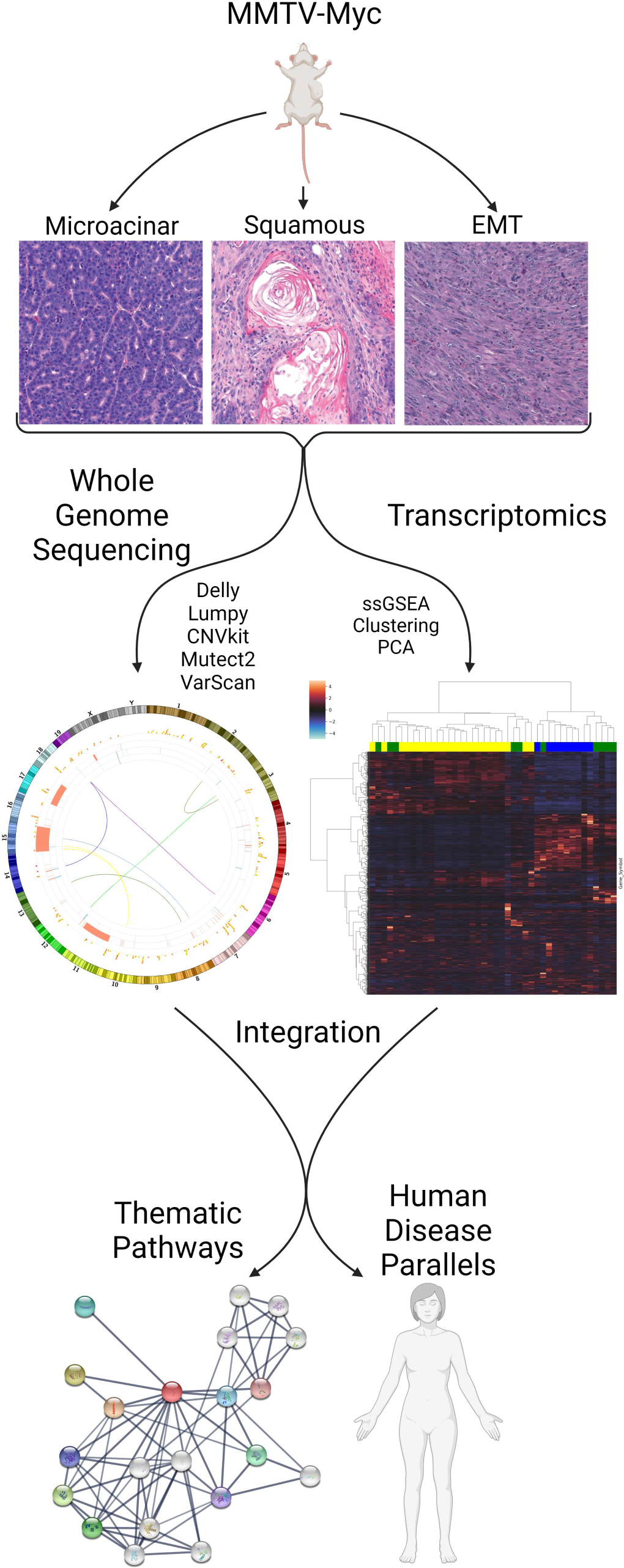
Schematic of workflow. Significant histological and transcriptional heterogeneity was found in the MMTV-Myc mouse model of human breast cancer. To ascertain the genetic origins of transcriptional differences and integrate these omics data between histological subtypes, we performed short read whole genome sequencing at a depth of 40x for three tumors of each unique and predominant histological subtype: microacinar, squamous, and EMT like tumors. Somatic single nucleotide variants, copy number alterations, inversions, and translocations were profiled after alignment and processing of genomic data. Genomic somatic variants were integrated with and compared to previously obtained Affymetrix microarray gene expression and pathway analysis by single sample gene set enrichment analysis (ssGSEA). Integration of multi-omic data enables the identification of thematic pathways driving tumorigenesis and putative oncogenic drivers for each histological subtype. Importantly, these thematic pathways identified in the MMTV-Myc mouse model share similarities in human breast cancer, impactful analogous genetic events in humans co-occur with Myc amplification, and these pathways affect human breast cancer patient overall survival significantly (Created with BioRender.com).

**Figure 2:**
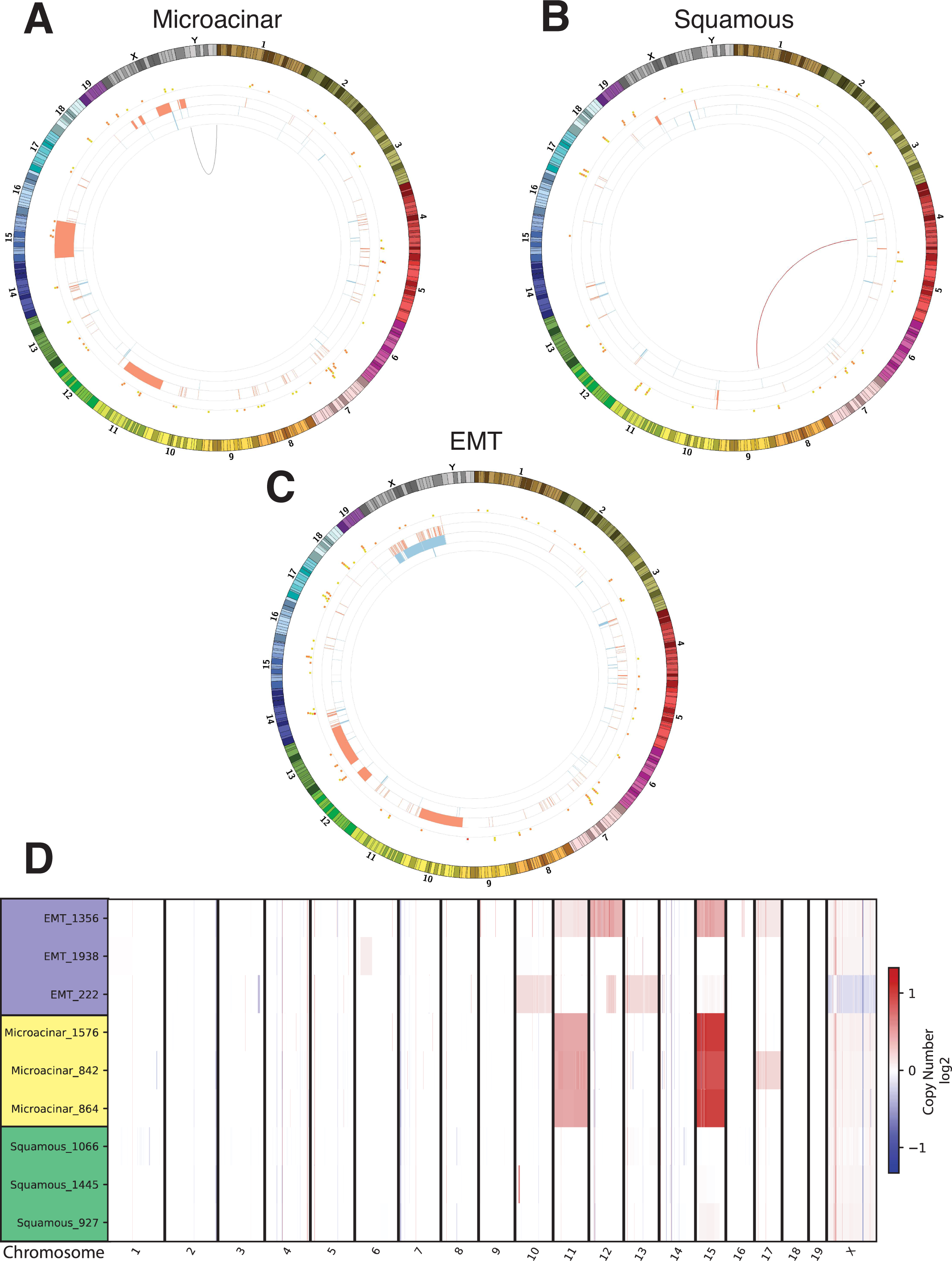
Heterogenous and conserved somatic features revealed by whole genome sequencing. Representative Circos plots are shown for (**A**) microacinar, (**B**) squamous, and (**C**) EMT histological subtypes. The outermost ring of each Circos plot depicts an ideogram for the mouse chromosomes proportionate with actual chromosome length. The next inner ring shows mutations in genes as stacked blocks at their corresponding genomic locations, color coded to their predicted impact by SnpEff^77^—yellow for low impact, orange for moderate impact, and red for high impact. The next inner ring shows discrete copy number changes as analyzed by CNVKIT; red regions indicate amplification and blue regions indicate deletions. The height of each copy number alteration corresponds to the predicted change in copy number, with the lowest level change being +-1 and showing a max copy number change of +-2. The innermost ring reveals inversions and translocations as determined by the consensus of Delly and Lumpy somatic variant callers. Inversions are colored black while translocations match the color of the ideogram of one of the two chromosomes involved in the translocation event. (**D**) A CNVKIT heatmap shows the log_2_ fold change of the estimated normalized copy number segments of each chromosome for each tumor sample relative to the wildtype reference.

Strikingly, all microacinar tumors sequenced shared the same whole chromosome amplification events on chromosomes 15 and 11 as revealed by copy number segmentation calls (**Figure 2D**). Estimated total ploidy gain for chromosome 15 is 2, for a total of 4 gene copies, and ploidy gain of 1 for chromosome 11, for a total of 3 gene copies. The predicted integer copy number gains were consistent across all microacinar tumors. While many whole chromosome amplifications and deletions were observed in EMT tumors, none were consistent across the histological subtype. Interestingly, there were very few focal and whole chromosome CNVs in individual squamous tumors and none shared across the subtype. The majority of CNVs across all tumors sequenced were broad and affected the entire chromosome rather than a discrete region within the chromosome, despite notable exceptions discussed in later figures. This is consistent with human breast cancer CNVs which often affect a whole chromosome arm on either side of the centromere^32^ except in the cases of strong selective pressure over a particular region, such as the cases of ERBB2 in breast cancer^33^ or AR in prostate cancer^34^.

### Copy number alterations are largely responsible for gene expression changes

It is well established that CNVs typically correlate strongly with changes in gene expression across cancer types^35^ and have been used to target cancer dependencies for therapeutic intervention^36^. Despite this, we sought to establish correlation between gene expression and CNVs in the MMTV-Myc mouse model tumors empirically. When correlating the estimated absolute integer copy number gain or loss for each gene with their matched gene expression sample and extending these results to 33 additional samples profiled by microarray, we obtained high Pearson’s *r* and Kendall’s *τ* correlation coefficients broadly across the genome for CNV sites (**Figure 3A**). Of note, the highest and most consistent correlation for both Pearson’s *r* and Kendall’s *τ* occurs on chromosomes 15 and 11, suggesting the highly conserved ploidy gains across microacinar samples translates well into increased gene expression.

**Figure 3:**
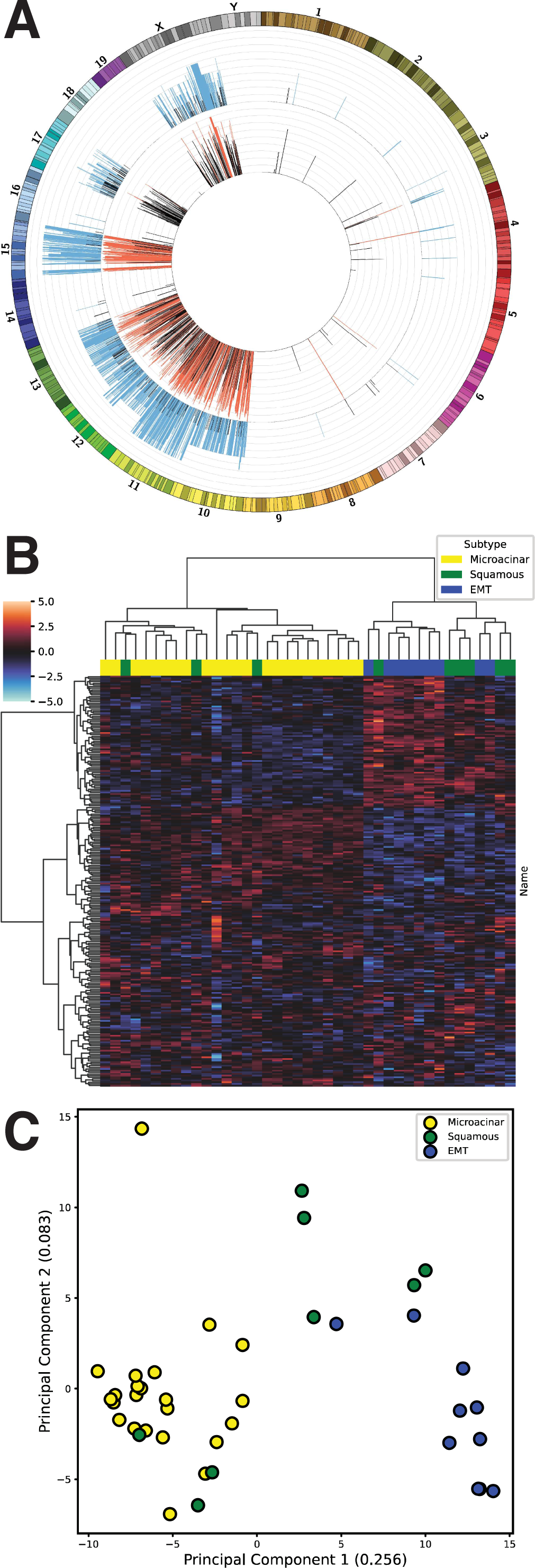
Copy number changes drive gene expression changes. (**A**) Circos plot of correlation of copy number changes and gene expression across all tumors samples. Significant Kendall’s rank correlation coefficient (>= 0.3) shown in blue and significant Pearson correlation coefficient (>= 0.7) shown in red. (**B**) Unsupervised hierarchical clustering of the MSigDB C1 positional dataset for ssGSEA values closely recapitulates the stratification of histological subtypes by gene expression clustering. (**C**) Principal component analysis (PCA) of C1 positional ssGSEA values reveals distinct clustering by histological subtype.

After establishing the link between CNVs and gene expression in the histological subtypes, we wanted to examine whether gene expression differences localized to human defined cytogenetic bands could stratify the histological subtypes. Indeed, when performing ssGSEA on gene expression data using the MSigDB C1 positional gene set, we find that unsupervised hierarchical clustering of these data post normalization largely clusters histological subtypes separately (**Figure 3B**), recapitulating clustering from raw gene expression data^13^. These data suggest CNVs or other events confined to specific genomic regions are largely responsible for gene expression differences between histological subtypes.

Principal component analysis (PCA) on these same C1 ssGSEA values reveals a similar clustering pattern among subtypes (**Figure 3C**). Principal component 1 was able to explain over 25% of the variance in this population, which largely separated the EMT and microacinar subtypes, with squamous tumors infiltrating both clusters or being separated mostly by principal component 2. In determining the number of clusters represented within this population, we employed affinity propagation^37^ on PCA initialized and non-PCA initialized C1 ssGSEA data with preferences set to the input similarities median value. The estimated number of clusters is 5 for PCA initialized data (**Supplementary Fig. 1**) and 6 clusters for non-PCA initialized data (**Supplementary Fig. 2**), possibly owing to further sub-heterogeneity within these populations.

### Differential mutational landscapes between mouse models of breast cancer

Having established CNVs as being a critical factor in determining gene expression in MMTV-Myc tumors, we sought to investigate whether differing mutations within these tumors played a role in gene expression changes. After limiting gene expression data to genes predicted to have moderate or high impact mutations, we did not see a statistically significant difference in the proportion of genes that were differentially regulated compared to the overall proportion of all genes that are differentially regulated between subtypes (data not shown). This is unsurprising as mutations typically do not affect self-gene expression, except in the case of truncating mutations^38^.

Next we examined wholistic mutation patterns between each histological subtype and compare them to mutation patterns in previously sequenced MMTV-Neu and MMTV-PyMT tumors^27^. We found no large differences in overall mutational burden between MMTV-Myc subtypes, although we found trends within each mouse model (**Figure 4A**). Mutational burden did not diverge significantly between MMTV-Myc subtypes but varied considerably in the MMTV-Neu mouse model. The MMTV-PyMT model exhibited the highest overall mutational burden of mouse models analyzed, but also demonstrated considerable variability between individual tumors.

**Figure 4:**
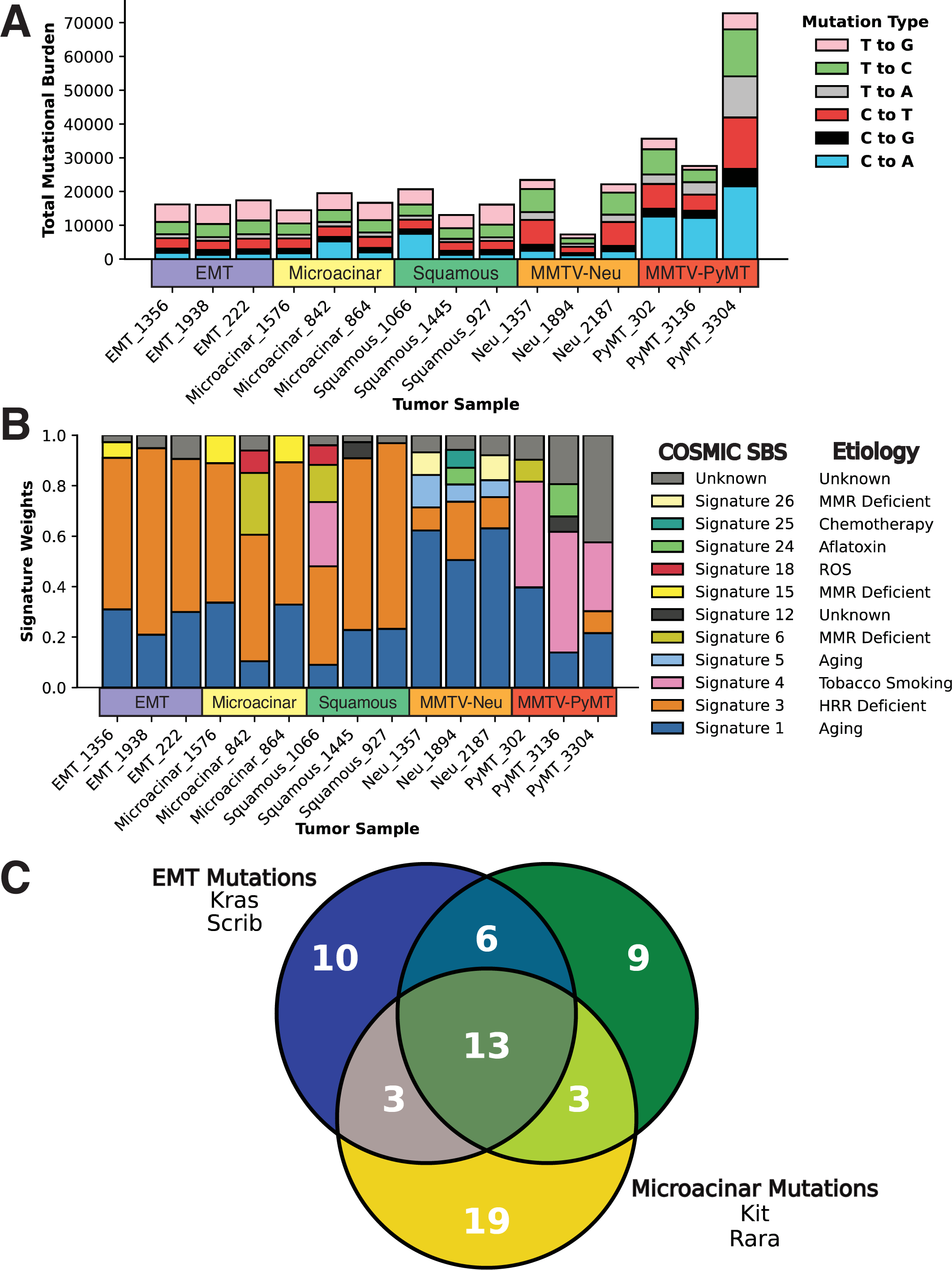
Oncogenic drivers determine mutational heterogeneity. (**A**) Total counts of overall somatic mutational burden of MMTV-Myc tumors compared to mutational burden of MMTV-Neu and MMTV-PyMT tumors as shown by bar plot. (**B**) Weights of Catalogue of Somatic Mutations in Cancer (COSMIC) mutational signatures derived from each tumor using DeconstructSigs^41^ depicted by stacked bar plot. (**C**) Venn diagram of representative conserved mutations (≥66% of tumors) between histological subtypes of moderate or high impact predicted by SnpEff.

We hypothesized these differences in mutational burden could arise from separate oncogenic drivers and mutational processes within each mouse model given the extensive characterization of mutational processes in human cancers^39,40^. To examine mutational processes in our mouse models, we utilized deconstructSigs^41^ to generate different COSMIC single base substitution (SBS) signatures that take adjacent nucleotides into context and ascribes etiologies to different trinucleotide mutational patterns. We found that all MMTV-Myc tumors regardless of subtype were predominated by the homologous recombination deficient (HRR) signature (**Figure 4B**). The HRR signature is strongly associated with germline and somatic mutations in BRCA1 and BRCA2 mutations in human breast cancer^39^. No mutations in BRCA1 or BRCA2 were found in any of the MMTV-Myc tumors analyzed, with these signatures possibly the result of promoter hypermethylation or other factors not analyzed. MMTV-Neu tumors were predominated by the clock-like aging signatures, while MMTV-PyMT tumors demonstrated large tobacco smoking signatures. Given that all mouse models were raised in similar controlled environments, these data suggest an unidentified endogenous C>A mutational mechanism present in MMTV-PyMT tumors.

While overall mutational burden and mutational signatures cannot parse between histological subtypes of the MMTV-Myc tumors, we reasoned that specific mutations may be associated with each subtype. Indeed, we find a small number of conserved (≥66% of tumors in each subtype) and impactful mutations within each subtype (**Figure 4C**). Notable conserved mutations in the EMT subtype include KRAS G12D activating mutations and splice variants in SCRIB. This may be significant as others have found SCRIB cooperates with MYC for transformation and mislocalization of SCRIB within the cell, sufficient to promote cell transformation^42^. Interestingly, the squamous subtype had no discernible conserved and impactful mutations between tumors. There may be heterogeneous conservation of signaling pathways activated at the transcriptional level, though, as each squamous tumor had impactful mutations in transcription factors: zinc finger and BTB domain containing (ZBTB) genes, zinc-finger protein (ZFP) genes, or both. For the microacinar tumors, we observed conserved missense mutations at A538E in proto-oncogene c-KIT (KIT) and at A255D for retinoic acid receptor-α (RARA). KIT is a well-established oncogene, particularly in acute myeloid leukemia (AML)^43^ where mutations play a large role in pathology, and RARA is involved in embryonic development whose disruption is well established in carcinogenesis^44^.

KIT and RARA missense mutations were found to be co-occurring with 100% frequency in 5 of the 9 total tumors sequenced, with no mutations in either gene standalone in the remaining 4 tumors. To validate our WGS findings and evaluate the extent of these mutations further in other MMTV-Myc tumors, we performed sanger sequencing on the tumors that previously underwent WGS and an additional two tumors from each histological subtype. From these data, we confirmed that these KIT and RARA mutations were co-occurring with 100% frequency in 7 of the 15 tumors that underwent sanger sequencing. From the total populations analyzed, these mutations were present in 60% of both the microacinar and squamous subtypes, while only 20% in EMT (**Supplementary Files 1 & 2**). These data may suggest a link between KIT and RARA given their perfect co-occurrence patterns, but the functional implications of these mutations are unknown at this time.

### Heterogeneous activation of KRAS signaling in the EMT subtype

Previous studies on the MMTV-Myc model have shown that the EMT subtype often possessed activating KRAS mutations and increased RAS signaling^13^, whereas the squamous and microacinar subtypes largely did not (**Figure 5A**). This was the case for EMT tumors 222 and 1938 that underwent WGS, where both had KRAS G12D missense mutations, while the squamous tumor 1066 acquired the less transforming KRAS G13R mutation (**Figure 5B**). Consequently, when comparing ssGSEA values for the C6 oncogenic signature gene sets between the three histological subtypes, EMT tumors consistently had upregulation of various KRAS signaling pathways across tissue types (**Figure 5C**). Investigation of a representative KRAS signaling pathway shows the probability of KRAS activation is considerably higher in EMT than both microacinar and squamous samples (**Figure 5D**).

**Figure 5:**
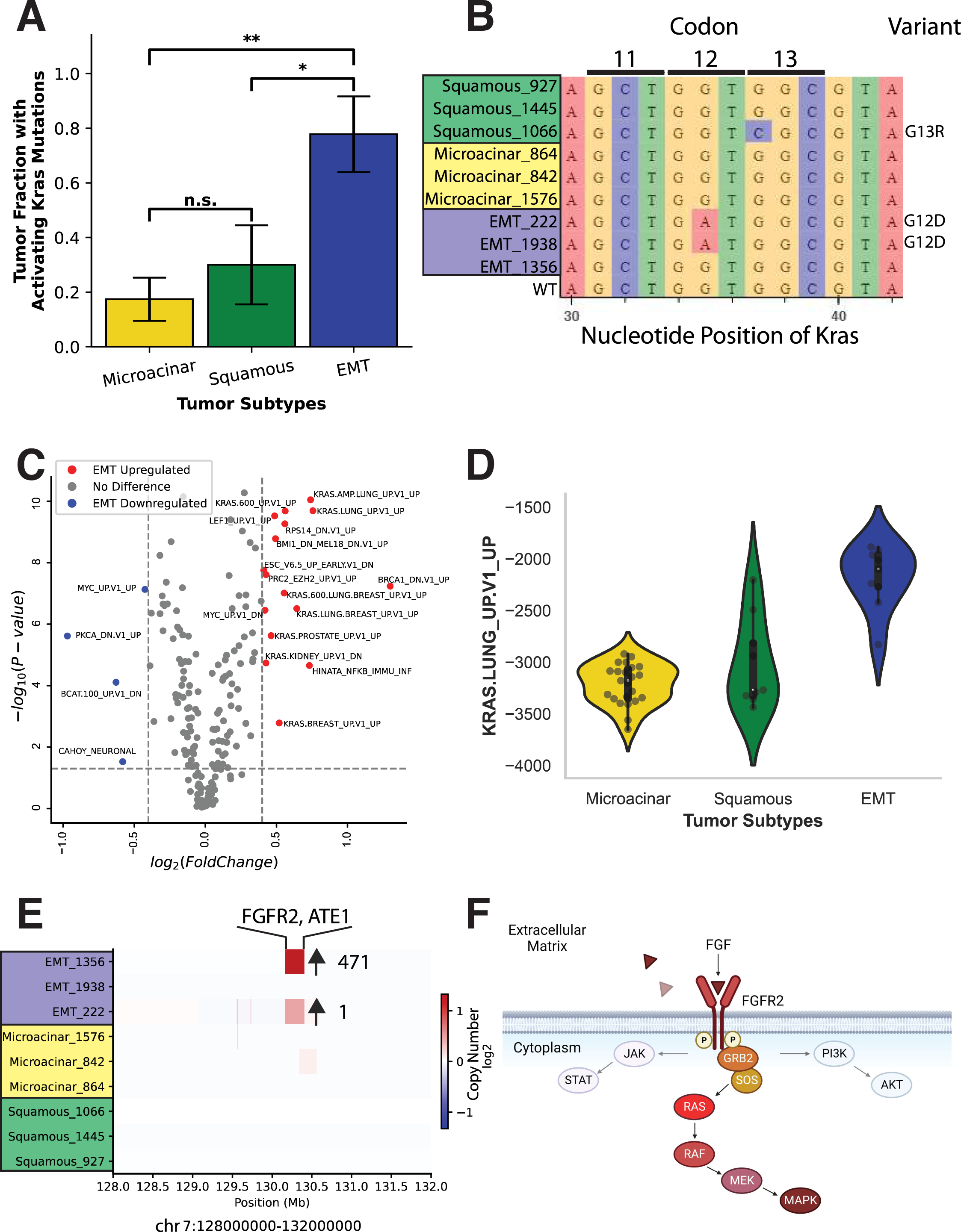
Heterogenous activation of KRAS pathway in EMT histological subtype. (**A**) Proportion of each tumor histological subtype with activating mutations in KRAS in bar plot format. (**B**) Sequence variation in KRAS between all tumors sequenced as shown by a logo plot illustrates canonically activating mutations in KRAS. (**C**) A volcano plot of the ssGSEA values from the MSigDB C6 oncogenic signature gene set, showing EMT upregulated or downregulated gene sets compared to microacinar and squamous. (**D**) Violin plot of a representative KRAS pathway signature from the ssGSEA values of the C6 oncogenic signature gene set, showing distinct upregulation of KRAS signaling in the EMT subtype. (**E**) Heatmap of log2 fold change in copy number segmentation values showing high level amplification of FGFR2 in EMT. (**F**) Canonical molecular pathway signaling reveals FGFR2 lies directly upstream of KRAS (Created with BioRender.com).

Despite its high ssGSEA KRAS activity scores **(Supplementary Table 1**), EMT tumor 1356 possessed no KRAS mutations or mutations in genes elsewhere in the RAS pathway. Copy number segmentation data obtained pointed to a high-level amplification event on chromosome 7, encompassing FGFR2 and ATE1 (**Figure 5E**). Of particular significance is the estimated copy number gain of this region. EMT tumor 222 had a similarly bounded focal amplification event over FGFR2 and ATE1 that increased both gene copy numbers by 1, but EMT tumor 1356 has an estimated copy number gain of 471 over this region and correlates well with gene expression levels of FGFR2 (**Supplementary Tables 2 & 3**). The scale of this amplification event suggests a strong selective pressure for focal amplification of FGFR2 in EMT tumor 1356. When examining the pathways of FGFR2 in the literature, FGFR2 acts directly upstream of RAS-MAPK, PI3K-AKT, and JAK-STAT signaling pathways^45^ (**Figure 5F**). Thus, we propose that the increase in KRAS signaling seen in EMT tumor 1356 is due to the large, focal amplification of FGFR2 and subsequent activation of RAS signaling. Stemming from the previous assessment, it is likely that the EMT histological phenotype is dependent on increased RAS signaling, ostensibly through heterogeneous genetic mechanisms.

### Squamous tumors represent an intermediate phenotype between microacinar and EMT

To this point, there are strong associations between copy number amplifications specific to the microacinar subtype and heterogeneous genetic events that lead to KRAS pathway activation in the EMT subtype. However, there are no readily identifiable conserved genetic features that can explain the squamous phenotype. Despite this, squamous tumors consistently occupy a gene expression state between that of the EMT and microacinar subtypes. PCA of C2 curated gene sets (**Figure 6A**) and C6 oncogenic signature gene sets (**Figure 6B**) show clustering of squamous samples between microacinar and EMT, with some squamous samples invading both the microacinar and EMT clusters. Affinity propagation estimates 3 clusters in PCA initialized and 4 clusters in uninitialized C2 ssGSEA signatures (**Supplementary Figs. 3 & 4**). Clusters for the C6 ssGSEA signatures are similar but slightly more heterogeneous, with 5 clusters estimated for both PCA initialized and non-initialized signatures (**Supplementary Figs. 5 & 6**).

**Figure 6:**
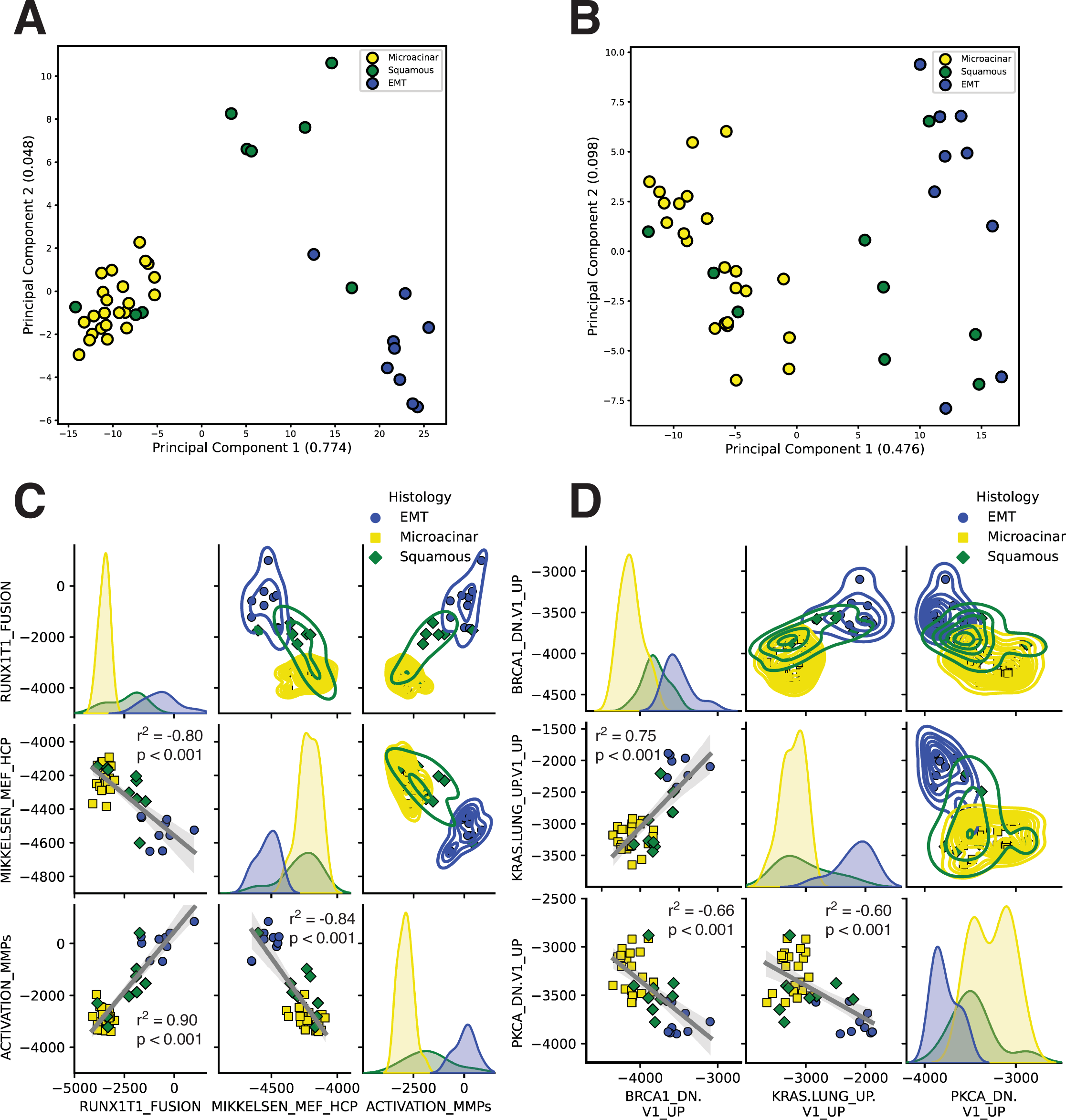
Squamous represents an intermediate phenotype between microacinar and EMT. (**A**) PCA of the MSigDB C2 curated and (**B**) C6 oncogenic signature gene sets for ssGSEA values of the microacinar, squamous, and EMT tumors recapitulates C1 clustering and explains more variance in the data. (**C**) Pairwise relationship plots for representative C2 gene set and (**D**) C6 gene set ssGSEA values are shown with linear regression lines and a 95% CI. Pearson R correlation values are shown with p-values determined from a two-sided t-test.

From the previous results, we sought to examine the relationship between individual pathways in each gene set that could explain these differences. To accomplish this, we chose the top three differentially expressed representative pathways from EMT and microacinar pathway signatures for C2 (**Figure 6C**) and C6 (**Figure 6D**) MSigDB gene sets. While there were no statistically significant correlations within each histological subtype (**data not shown**), likely due to limited numbers of samples, pairwise correlations of pathway signatures across all three subtypes showed both significant positive and negative correlations between ssGSEA pathway activities. Beyond this, top differentially expressed pathway activities appear to lie on a continuum, with squamous tumors routinely spanning between and infiltrating the microacinar and EMT clusters. Pairwise relationships for the C1 positional gene sets shows similar patterns (**Supplementary Fig. 7**). These data are consistent with the squamous histology occupying an intermediate phenotype between that of EMT and microacinar subtypes.

### Integrated mouse data stratifies human breast cancer subtypes and yields clinical insights

It is clear that the MMTV-Myc mouse model produces primary mammary tumors that are heterogeneous in histology, gene expression, metastatic variance^13^, and now somatic genomic perturbations that can explain many of the transcriptional differences seen in this model. However, the significance of these events and translational potential to humans is not immediately obvious.

To evaluate whether the events seen in MMTV-Myc mouse model could have predictive power in clinical outcomes, we utilized an integrative approach, combining gene expression data and somatic genetic events to examine how they can parse human breast cancer subtypes. To this end, we took a subset of genes that were differentially expressed between MMTV-Myc histological subtypes and were also included in conserved copy number gain or copy number loss events within the model. Subsequently, we performed unsupervised hierarchical clustering on human gene expression from the METABRIC breast cancer dataset^46–48^ limited to the integrative mouse gene set (**Figure 7A**). We found distinct clusters emerged that well represented the intrinsic subtypes of breast cancer. Importantly, clustering from the same number of genes used but instead randomly selected failed to resolve intrinsic breast cancer subtype clusters to the same degree (**Supplementary Figs. 8-17**). This suggests that the integrated mouse gene set represents a diverse set of informative genes across all intrinsic subtypes of human breast cancer that can effectively differentiate between subtypes rather than contributing noise. Clustering by the PAM50 gene set showed improved subtype clustering overall relative to the mouse integrative gene set but did not resolve the luminal A and luminal B subtypes to the same extent (**Figure 7B**). It is important to note that the PAM50 gene set is curated specifically to be able to differentiate between human intrinsic breast cancer subtypes, so seeing improved performance relative to the integrative mouse gene set is expected.

**Figure 7:**
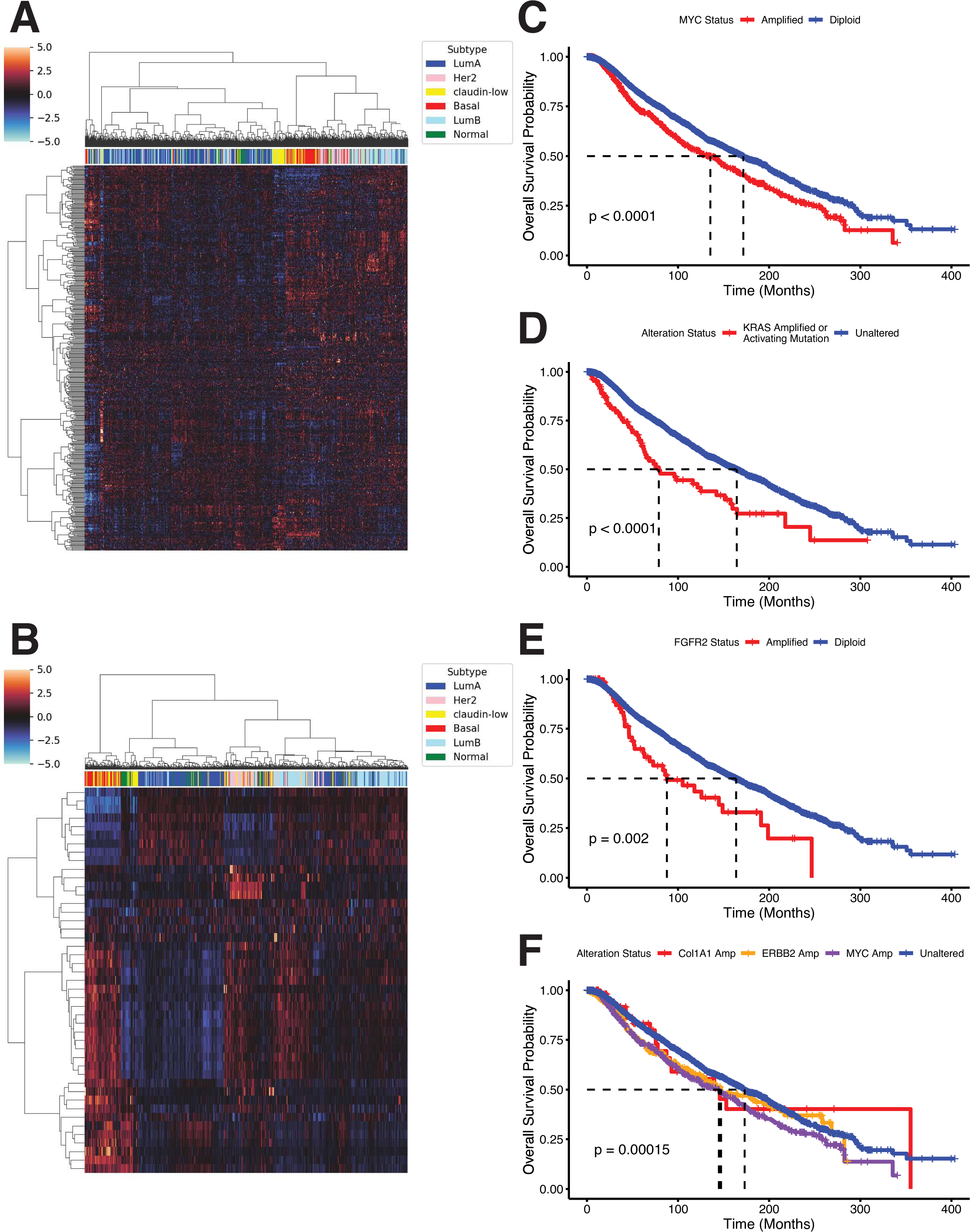
Mouse model genetic and transcriptional events inform human clinical outcomes. (**A**) Unsupervised hierarchical clustering of METABRIC gene expression values by a list of 453 homologous genes that are in a conserved amplification/deletion event and are differentially expressed between MMTV-Myc histological subtypes. (**B**) Unsupervised hierarchical clustering of METABRIC gene expression values by the PAM50 gene set. (**C**) Overall survival (OS) Kaplan Meier (KM) analysis of non-redundant TCGA breast cancer patients, accessed through cBioPortal, stratified by Myc amplification status. (**D**) OS KM curve of non-redundant TCGA breast cancer patients stratified by KRAS alteration status. (**E**) OS KM curve of non-redundant TCGA breast cancer patients stratified by FGFR2 amplification status. (**F**) OS KM curve of non-redundant TCGA breast cancer patients stratified by copy number alteration status to Col1A1, RNF43, or neither; overlapping samples, such as those with both Col1A1 and RNF43 amplification, are excluded from analysis.

**Figure 8:**
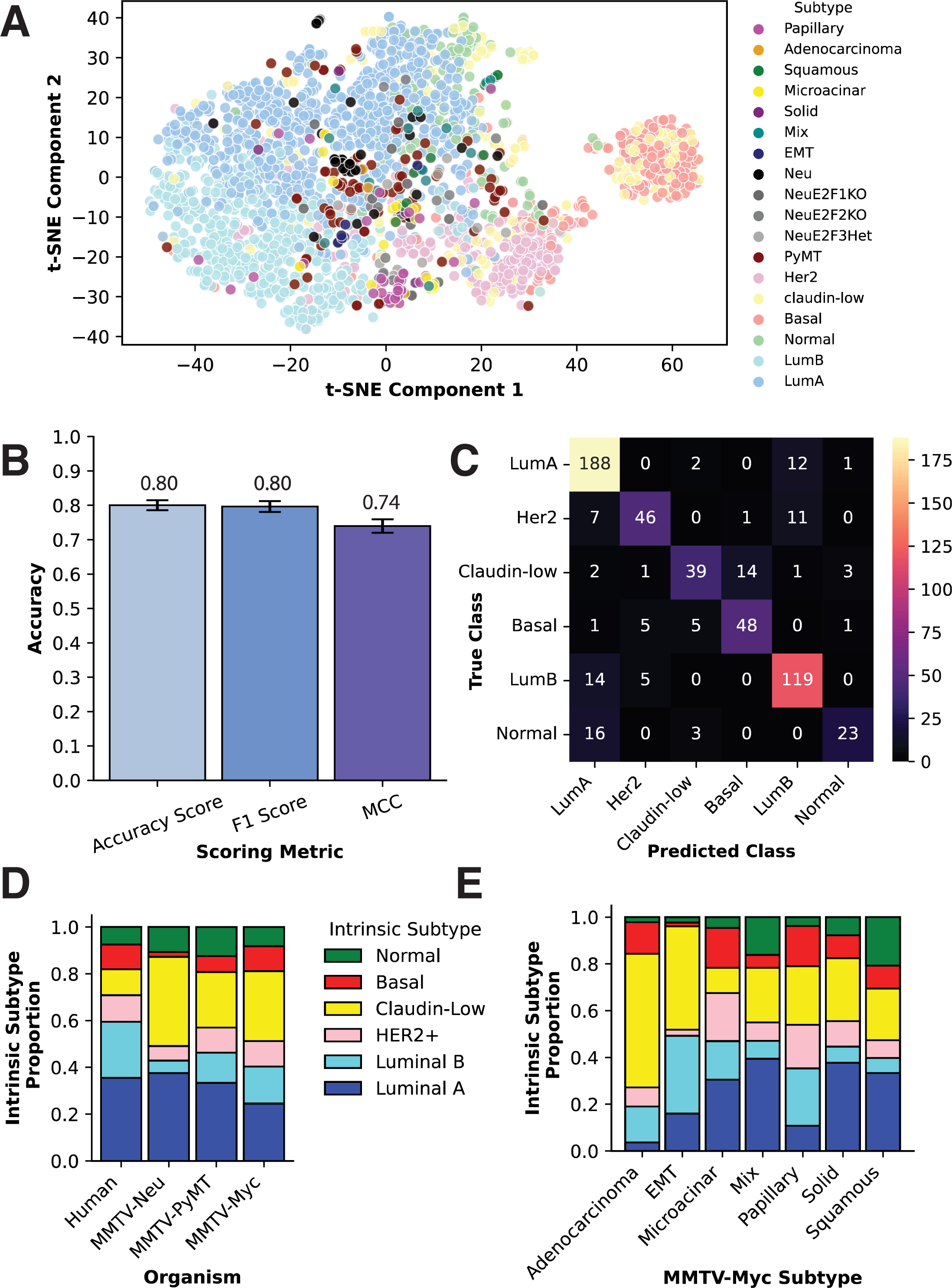
MMTV-Myc histological subtypes are representative of different human breast cancer intrinsic subtypes. (**A**) t-distributed stochastic neighbor embedding (t-SNE) performed on human METABRIC and mouse model gene expression samples using the 32 homologous gene subset of PAM50 as determined by recursive feature elimination with cross validation. (**B**) Normalized scoring metrics for the soft voting classifier including accuracy for measuring true positives, a weighted F1 scoring metric for balancing precision and recall, and a Matthews correlation coefficient (MCC) metric for taking into account false positives and false negatives even in the case of unbalanced classes. (**C**) A confusion matrix where true positives lie along the diagonal from top left to bottom right and false values occupy all other boxes. (**D**) Bar chart of proportional probabilities of each model representing human intrinsic breast cancer subtypes as determined by the soft voting classifier. Human intrinsic subtype proportion was determined directly from proportions of METABRIC breast cancer patients subtyped. (**E**) Bar chart of MMTV-Myc histological subtypes and proportional probabilities of each subtype corresponding to different human breast cancer intrinsic subtypes determined by the soft voting classifier.

To compare the effectiveness of this gene set in distinguishing clusters of human breast cancer patients quantitatively, affinity propagation was performed on non-PCA initialized METABRIC gene expression data over all genes, genes limited to PAM50, the integrated mouse gene set, and randomly selected gene sets corresponding to the same number of genes as the PAM50 and integrated mouse gene sets to serve as controls. The full METABRIC dataset converged on 1,886 clusters, possibly owing to the significant amount of noise in the bulk transcriptional data (**Supplementary Fig. 18**). Under the same conditions, the PAM50 gene set converged to 7 different clusters (**Supplementary Fig. 19**) and the integrated mouse gene set converged to 12 clusters (**Supplementary Fig. 20**). The random gene sets produced 3 clusters for 50 random genes (**Supplementary Fig. 21**) and 9 clusters for 446 random genes (**Supplementary Fig. 22**). However, the clusters generated for the PAM50 and integrated mouse gene set are well dispersed throughout the space compared to the random gene set clusters. This is significant as this shows both the PAM50 and integrated mouse gene sets are better able to resolve the intrinsic clusters of these breast cancer patients compared to random gene sets. These results are corroborated by PCA analysis on these same gene subsets where more variance can be explained by principal components of the PAM50 and integrated mouse gene set than random gene subsets (**Supplementary Figs, 23-27**). Separation of the luminal A, luminal B, and normal intrinsic subtypes is evident in the integrative mouse gene set compared to random gene sets or all genes.

It is clear that genetic events, particularly CNVs, in the mouse model can explain gene expression differences between each histological subtype and can resolve human breast cancer intrinsic subtypes. However, it is unclear to what extent these genetic events in the MMTV-Myc mouse model represent genetic events occurring in human breast cancer. To address this, we assayed publicly available TCGA datasets on breast cancer available through cBioPortal. We assessed prevalence of genetic events in human breast cancer first identified to be conserved both in genomic alteration status and differential gene expression in the MMTV-Myc mouse model. Subsequently, we examined the effects of these genes on human breast cancer overall survival clinical outcomes through Kaplan-Meier analysis.

All of the genetic events examined were found to be co-occurring with MYC amplification in human breast cancer (**Supplementary Table 4**), supporting the use of the MMTV-Myc mouse model in studying MYC driven human breast cancers. However, because of the co-occurring nature of these events and the limited number of patient samples with genetic events and matched clinical outcomes, statistical significance is difficult to achieve in mutually exclusive populations of MYC amplification and identified genetic events. MYC amplification and overexpression is well described in TNBC and basal like subtypes of breast cancer^1,49^, known as the deadliest subtypes of breast cancer currently^50,51^, which must be accounted for in survival analysis. To remedy this, we look at relative differences in overall survival from MYC amplification compared to analogous genetic events in humans identified in the mouse model. Indeed, we find steep decreases in overall survival in patients with analogous genetic events seen in MMTV-Myc tumors when compared to MYC amplification alone.

Kaplan-Meier analysis on breast cancer patients revealed MYC amplification was present in 18.2% of patients surveyed, with median overall survival at 135.2-months (95% CI: 113.7 – 148.1) compared to the unaltered population with median overall survival at 171.3-months (95% CI: 161.2 – 182.9) (**Figure 7C, Supplementary Tables 5 & 6**). When comparing the patient population of those harboring KRAS activating mutations or amplifications, which account for 2.2% of all breast cancer cases, we find a marked decrease in overall survival compared to the unaltered cohort (**Figure 7D**) with median overall survival at 77.7-months (95% CI: 61.8 – 146.4) for the KRAS altered population compared to median overall survival of 164.3-months (95% CI: 154.3 – 173.0) for the unaltered cohort (**Supplementary Tables 7 & 8)**. While it cannot be ruled out that MYC amplification co-occurrence in the KRAS altered cohort reduces overall survival, the KRAS altered population maintains a 57.5-month median overall survival deficit to MYC amplification alone. Thus, MCY amplification may only have a modest additive effect to KRAS amplification or activating mutations in this cohort. It is worth noting that the unaltered population in the KRAS cohort has reduced overall survival compared to the MYC unaltered cohort, suggesting most patients with MYC amplification are largely shunted to the unaltered group.

Similarly, patients with focal FGFR2 amplifications, accounting for 1.5% of all breast cancer cohort patients, also exhibit a marked decrease in overall survival compared to the unaltered cohort (**Figure 7E**) with median overall survival at 87.7-months (95% CI: 62.4 – 191.0), contrasting with the unaltered cohort at 163.5-months (95% CI: 154.0 – 172.9) (**Supplementary Tables 9 & 10**). Again, patients with FGFR2 amplification fare significantly worse than those with MYC amplification, standing at a difference of 47.5 months median overall survival difference. While KRAS and FGFR2 alterations are infrequent in breast cancer, these alterations may be extremely significant in the prognosis and treatment of their disease.

Interestingly, we observed conserved amplification in the microacinar subtype over mouse chromosome 11, containing Col1A1 and CHAD, previously identified as playing a role in increased migration and metastasis in breast cancer^27^. Mouse chromosome 11 also contains ERBB2; it is interesting to note that MYC and ERBB2 amplification often co-occur in human breast cancer, investigated by others previously^52,53^. To try and rule out secondary effects from ERBB2 and MYC amplification in patients with Col1A1 amplification, we stratified this breast cancer cohort into four categories: Col1A1 amplification exclusively (1.7% of patients), ERBB2 amplification exclusively (11.1% of patients), MYC amplification exclusively (17.3% of patients), and unaltered (remaining 69.9% of patients). This analysis reveals no significant differences between the Col1A1, ERBB2, and MYC amplified populations median overall survival (139.6-months, 146.8-months, and 144.2-months, respectively) (**Figure 7F, Supplementary Tables 11 & 12**). While there are too few Col1A1 exclusively amplified samples in this population to reach significance compared to the unaltered population by a pairwise log-rank test, when comparing non-exclusive Col1A1 amplified patients to the unaltered cohort we maintain a similar median overall survival at 142.4-months (p-value: 0.0041, **Supplementary Table 11**). These data show that the Col1A1 amplification event significantly decreases overall survival in breast cancer patients independently of ERBB2 and MYC amplified populations.

OBSCN was identified with conserved mutations across all histological subtypes, with OBSCN mutations correlating poorly with overall survival in humans (**Supplementary Fig. 28, Tables 13 & 14**). However, OBSCN is a large gene where mutations often co-occurr with other large genes, such as TTN, MUC family genes, and FAT family genes (**Supplementary Table 15**). Therefore, mutations in OBSCN are likely biomarkers of hypermutational burden rather than OBSCN playing a tumor suppressive or oncogenic role. This could still be useful information, as hypermutant individuals are more likely to respond to immune checkpoint inhibitors^54^.

Altogether, these analyses reveal conserved genetic events between both human Myc driven breast cancer and the MMTV-Myc mouse model of breast cancer. Importantly, these somatic genetic events are associated with severe drops in overall survival in human breast cancer patients in large excess of what Myc amplification causes.

### Machine learning classifier predicts MMTV-Myc histological subtypes correspond to different human breast cancer intrinsic subtypes

Thus far, we have shown that there exists tightly linked transcriptional and genomic perturbations in the MMTV-Myc model which are heterogeneous across its histological subtypes with clinical implications in humans. However, whether these histological subtypes are representative of different human breast cancer intrinsic subtypes is unclear. Unsupervised clustering by the integrative mouse gene set shows modest ability to parse human intrinsic breast cancer subtypes, but it falls short of being able to discriminate between features (genes) that can best exemplify a given class (intrinsic subtype). To this end, we employed a machine learning model classifier trained on human METABRIC gene expression data to predict which human breast cancer intrinsic subtype each MMTV-Myc histological subtype best represents.

To accomplish this, we first combined the raw METABRIC microarray gene expression data with the MMTV-Myc microarray data (GSE15904), along with additional mouse microarray cohorts for variations of the MMTV-Neu mouse model (GSE42533) and the MMTV-PyMT mouse model (GSE104397) for comparison. PCA of normalized data revealed strong batch effects between datasets that would distort causal biological interpretation (**Supplementary Fig. 29**). In removing batch effects, we employed the parametric empirical Bayes shrinkage adjustment available from ComBat^55^, effectively eliminating non-biological differences between datasets (**Supplementary Fig. 30**).

We then sought to narrow down the number of features in this dataset to avoid overfitting the model and make it more generalizable to new patient data for intrinsic subtype prediction. The prediction analysis of microarray 50 (PAM50) is a well-established scoring metric of gene expression data for 50 genes to stratify breast cancer patients into intrinsic subtypes and offer clinical prognoses^56^. From here, we employed recursive feature elimination with cross validation (RFECV) with a support vector machine (SVM) radial basis function kernel estimator to determine the optimal features in the dataset. Surprisingly, 32 specific genes (**Supplementary Table 16**) from the PAM50 subset gave higher cross validation scores than utilizing all 50 genes (**Supplementary Fig. 31**). In testing various estimators for the machine learning classifier model limited to this 32 gene subset, we found a great deal of variability between models in hard classification predictions. To alleviate this, we utilized a soft voting classifier composed of logistic regression, SVM, random forest, XGBoost, and multi-layer perceptron estimators with some hyperparameter tuning—exact specifications are listed in the supplemental Jupyter Notebooks. Pooling multiple estimators together was done to reduce bias from any one particular estimator and to increase overall accuracy. Subsequently, the highest average probability of a given class will decide which intrinsic subtype a mouse tumor belongs to.

The motivation for developing a supervised machine learning model classifier is evident after examining the distribution of the combined human and mouse gene expression dataset through a scatterplot of t-distributed stochastic neighbor embedding^57^ (t-SNE) data in two-dimensional space (**Figure 8A**). There are distinct clusters of human breast cancer samples forming the intrinsic subtypes except for claudin-low, which supports the notion that claudin-low breast cancers represent an additional complex phenotype rather than an intrinsic subtype of breast cancer^58^. The mouse samples are well dispersed throughout the plot even within the same mouse model, pointing to substantial gene expression heterogeneity within each model. However, problems arise in that it is not immediately obvious which cluster each mouse sample belongs to. When using t-SNE, there is also necessarily a loss of information even when performing non-linear dimensionality reduction, in this case from 32 to 2 dimensions. This is nothing to say of the tendency for gradient descent algorithms such as t-SNE to become stuck in local optima. Additionally, other unsupervised methods such as hierarchical clustering equally weight all features, which decreases accuracy of the model. For these reasons and more, there was a clear need to develop a supervised machine learning classifier using the 32 gene dataset.

Accuracy scores and F1 scores are often used to evaluate the efficacy of supervised machine learning models. However, it has been shown that these metrics can be overinflated^59^, and so we also report the more robust Matthew’s Correlation Coefficient (MMC) metric which proportionally accounts for true positives, true negatives, false positives, and false negatives. For the soft voting classifier used to predict mouse-to-human subtypes, we obtained an average accuracy score of 80.0%, a weighted F1 score of 80.0%, and an MCC score of 74.0% after 15-fold stratified and shuffled cross validation using a 70% train and 30% test split (**Figure 8B**). All metrics used to evaluate the machine learning model are in high agreement and show the model is effective at predicting human breast cancer intrinsic subtypes. To better visualize these predictions on the test data, we constructed a confusion matrix showing true classes on the vertical axis and predicted class on the horizontal axis (**Figure 8C**). Many of the most confused classes correspond well to the t-SNE visualization where clusters overlap, including the overlap of luminal A and luminal B classes, overlap of luminal A and normal classes, and the overlap of claudin-low and basal classes. All metrics combined show that the initial METABRIC gene expression data limited to the 32 most predictive genes shared with the mouse microarray data has significant power in discriminating between intrinsic subtypes of breast cancer using the soft voting classifier.

When applying the classification model to mouse gene expression data, we obtain heterogeneous subtype predictions across the various MMTV-Myc, MMTV-Neu, and MMTV-PyMT based mouse models (**Figure 8D**). Our initial hypothesis postulated that the MMTV-Myc model would enrich for claudin-low and basal-like tumors, similar to how human breast cancer patients with amplified MYC are preferentially basal-like or claudin-low. Indeed, for MMTV-Myc tumors overall, 30% are predicted as claudin-low, 24% are luminal A, 16% are luminal B, 11% are basal, 11% are HER2+, and 8% are normal-like. Large differences in intrinsic subtype proportions are evident across all mouse models tested in comparison with human intrinsic breast cancer subtype proportions, with METABRIC intrinsic subtype proportions at 35% luminal A, 24% luminal B, 11% HER2+, 11% claudin-low, 11% basal, and 8% normal-like. The claudin-low subtype is especially enriched across all mouse models examined.

Though, ratios of each intrinsic subtype vary considerably between histological subtypes for the MMTV-Myc model (**Figure 8E**). Adenocarcinoma tumors are primarily categorized as claudin-low at 57%, with luminal B and basal subtypes trailing at 15% and 14%, respectively. Papillary tumors maintain roughly similar proportions of basal (17%), claudin-low (25%), HER2+ (19%), and luminal B (24%) subtypes. EMT tumors are overwhelmingly claudin-low (44%) and luminal B (33%). Microacinar tumors show the most enrichment for the HER2+ subtype at 21% and tied for most basal enriched with papillary at 17%. Squamous tumors are considerably variable, with 33% classified as luminal A, 22% as claudin-low, and 21% as normal-like, corroborating previous pathway signatures showing squamous tumors are not confined to a specific localized cluster.

These results have the potential to inform mouse model use when investigating different subtypes of breast cancer or examining breast cancer heterogeneity more generally. For instance, the MMTV-Neu mouse model is often used as a model of HER2+ breast cancer, but most MMTV-Neu tumors show gene expression similar to luminal A and claudin-low human tumors. While other groups have predicted MMTV-PyMT tumors to correspond to the luminal B subtype^19^, we predict only 13% of PyMT based mouse models fall into this category, with PyMT tumors overall being quite heterogeneous.

In summary, we have created an accurate supervised machine learning classification model that can stratify human breast cancer intrinsic subtypes. When applied to batch effect corrected mouse transcriptional data, we observe diverse intrinsic subtype profiles assigned to different histological subtypes from the MMTV-Myc mouse model.

## Discussion

The utility of mouse models of breast cancer has progressed from overexpression of a driving oncogene to questions as to whether they accurately mimic the heterogeneity and progression of human breast cancer. Here, we have described the utility of the MMTV-Myc GEMM in recapitulating the histological, transcriptional, and genomic heterogeneity of human breast cancer.

Conserved somatic genetic changes across MMTV-Myc histological subtypes are associated with negative effects on overall survival when applied to human breast cancer patients. These somatic events have largely been overlooked previously due to their low prevalence in human breast cancer and resulting lack of statistically significant differences in clinical outcomes. Given that MYC is frequently amplified in basal-like and TNBCs^1^, and the current lack of targeted therapies in TNBCs^60^, it is likely there is no consistent driver of oncogenesis across all TNBC patients. Instead, we hypothesize there may be low prevalence oncogenic drivers that lead to similar transcriptional profiles ultimately, but each patient’s tumor maintains different genetic drivers. An example that arises in this paper is that of activating KRAS mutations or significant ploidy gain of FGFR2. Activation of either proto-oncogene will result in increased mitogen-activated protein kinase (MAPK) signaling, but treating the root oncogenic driver will require different therapeutic strategies. The most prevalent KRAS mutations in breast cancer are G12C/D/V/A mutations, with some of these G12C mutant patients potentially benefitting from treatment with the recently developed sotorasib^61^ or adagrasib^62^ therapies. However, patients with amplified FGFR2 would not respond to these therapies, and instead would more likely benefit from highly selective FGFR inhibitors such as AZD4547. A recent clinical study found that treating endocrine therapy resistant breast cancer patients with AZD4547 achieved partial response in some patients, with differentially expressed genes involved in FGFR signaling able to distinguish between responders and non-responders^63^. Although FGFR1 is amplified more often in breast cancer, both FGFR1 and FGFR2 amplified and overexpressing breast cancers could likewise benefit from AZD4547 treatment.

Aside from direct clinical implications in humans, these results reveal human intrinsic subtype analogs in the MMTV-Myc, MMTV-Neu, and MMTV-PyMT mouse model histological subtypes. Others have classified mouse models of breast cancer and ascribed them to different human intrinsic subtypes of breast cancer previously in unsupervised methods^18,19^. However, these analyses are not weighted for genes that are able to discriminate between intrinsic subtypes, they may be biased towards noise in the dataset rather than predictive signals, and do not produce metrics for scoring accuracy of the model. To rectify this, we created an accurate machine learning classification model trained on human gene expression data and applied to batch effect corrected gene expression data of different mouse models of breast cancer. From these results we see there is substantial heterogeneity within the histological subtypes of the MMTV-Myc model.

However, the proportions in MMTV-Myc tumors that match human intrinsic subtypes are skewed relative to their occurrence in humans. We see a general decrease in luminal tumors and an increase of claudin-low and normal-like tumors. While this is true overall for the MMTV-Myc mouse model, it is highly dependent on the histology of the tumor. For example, adenocarcinoma tumors are 57% claudin-low while microacinar tumors are 11% claudin-low. EMT tumors are largely mapped to claudin-low and luminal B intrinsic subtypes. The EMT subtype maintains great variability in CNVs, similar to that of human luminal B breast cancer^64^, but the gene expression signatures of EMT tumors overlap with the canonical signatures associated with human claudin-low breast cancer: high expression of markers for cytotoxic T-cell and natural killer (NK) cell infiltration (Granzymes C, D, E, F, and G), high expression of dormancy markers (NR2F1), and low expression of cell-adhesion proteins (CLDN2, GJB1, CEACAM1)^65^ (**Supplementary Fig. 32**). These observations may be useful to cancer researchers when selecting a mouse model for studying a specific subtype of breast cancer, particularly the adenocarcinoma or EMT histological subtypes as there are no established spontaneous models of breast cancer that exclusively mimic claudin-low breast cancer^19^. This data suggests that the adenocarcinoma or EMT histological subtypes from the MMTV-Myc GEMM could be a reliable immunocompetent model for claudin-low breast cancer.

It is clear that the copy number changes identified among the MMTV-Myc tumors sequenced, particularly the conserved ploidy gains on chromosomes 11 and 15 in the microacinar tumors, correlate with gene expression changes. ERBB2 and related genes lie on chromosome 11, which may explain the propensity of microacinar tumors to be HER2+ like than other histological subtypes. It is likely these CNVs are causal in driving gene expression changes between histological subtypes, although we cannot confirm that with these limited data. These highly conserved copy number gains seen in the microacinar tumors suggest a strong selective pressure for amplification and overexpression of genes in these regions. Upon examining the human homologs and their syntenic chromosomal regions for the integrated mouse gene set, we find the two largest high synteny regions are the entirety of chromosome 17 following from the mouse chromosome 11 amplification and the long arm of chromosome 8 (8q) following from the mouse chromosome 15 amplification (**Supplementary Table 17**). It is interesting to note that human chromosome 8q contains the MYC locus and chromosome 17q contains the ERBB2 locus, with amplification of these two loci highly correlated in human breast cancer (**Supplementary Table 18**). Regions 8q and 17q are among the most common frequently amplified regions in human breast cancer^66,67^. However, given the small sample size of microacinar tumors used and experimental setup, it is impossible to determine if either of these amplifications has causal implications for the other. It should also be noted that chromosome 17p is often deleted in many human breast cancers while the entirety of chromosome 17 in humans is able to be mapped to mouse chromosome 11. Mouse chromosomes are telocentric and given that these frequent large amplifications lie on either side of the centromere in humans, it is possible that amplification of the analogous genes from region 17q is of higher selective consequence than deletion of the analogous genes from region 17p. Comparative genomics between the MMTV-Myc histological subtypes and MYC driven human breast cancers may be an important area of future study.

## Conclusions

A significant hurdle in the study of breast cancer *in vivo* has been the limitations of mouse models recapitulating the heterogeneity found in human breast cancer. Here we report that the MMTV-Myc GEMM recapitulates the histological, transcriptional, and genomic heterogeneity found in human breast cancer, with important clinical parallels identified. We find different MMTV-Myc histological subtypes preferentially represent different human intrinsic breast cancer subtypes, further solidifying the MMTV-Myc model as an appropriate *in vivo* method for examining the multi-faceted aspects of human breast cancer heterogeneity even down to the gene expression level.

## Methods

### Whole Genome Sequencing

Flash frozen MMTV-Myc tumors^13^ stored at −80 °C were ground using a sterile mortar and pestle under liquid nitrogen. 3 tumors each from the microacinar, squamous, and EMT MMTV-Myc histological subtypes underwent DNA extraction. DNA was extracted for each tumor using a Qiagen Genomic-tip 20/G KIT according to manufacturer specifications. Whole genome sequencing was performed at Michigan State University’s Research Technology Support Facility (RTSF). DNA was sequenced at 40x depth using Illumina HiSeq 2500 paired end 150 base pair reads after TruSeq Nano DNA library construction.

### Transcriptomics

Gene expression data used in this study has previously been published^13^ on MMTV-Myc and MMTV-Neu tumors and is publicly available in the Gene Expression Omnibus (GEO) under accession number GSE15904. MMTV-PyMT and MMTV-PyMT E2F knockout tumor transcriptional data has been published^68^ and is available under GEO accession number GSE104397. MMTV-Neu E2F knockout transcriptional data was previously published^69^ and is available under accession number GSE42533.

### Whole Genome Sequence Processing

Raw whole genome sequencing paired end fastq files first underwent initial quality control using FASTQC^70^. Quality and adapter trimming of reads was performed using Trimmomatic^71^, with reads reassessed for quality afterwards again using FASTQC. Reads were then aligned to the mm10 reference *Mus musculus* genome using BWA-mem^72^ with option -M selected for compatibility with Picard. After alignment, read groups were added using Picard^73^. SAMtools^74^ was then used to sort bam files, mark PCR duplicated sequences, and index bam files. Discordant and splitter read files were also generated and sorted using SAMtools for downstream somatic variant callers.

### Somatic Variant Calling

Somatic mutations were called using the consensus of Mutect2^75^ (GATK suite) and VarScan^76^ calls based on chromosome, position, reference base, and variant base. Variants were then annotated using SnpEff^77^. To reduce false positives, filtering included subtracting FVB specific mutations from the mm10 C57BL/6 background as indicated in the FVB_NJ.mgp.v5.snps.dbSNP142.vcf file from the Wellcome Sanger Institute. In addition to this, HaplotypeCaller on GATK was used to call germline mutations on the FVB_NJ genome REL-1604-BAM available from the Sanger Mouse Genomes Project^78^ FTP server. After converting the FVB_NJ.bam WT reference to fastq files using SAMtools, these fastq files underwent the same whole genome sequence processing as MMTV-Myc tumors. Germline variants from this wiltdtype FVB background against the mm10 reference genome were subtracted from somatic variants in each tumor.

Somatic copy number variations were determined using multiple methods; discrete copy number variations are based on the consensus of Delly^79^ and Lumpy^80^ calls, while whole chromosome ploidy count and segmentations were determined using CNVKIT^81^. For discrete CNVs, only those with length above 10,000 base pairs, no evidence in the WT background, and a mapping quality (MAPQ) score at 60 or above were included for analysis. Delly and Lumpy calls were combined by similar genome starting and ending positions within a 100 base pair margin of error for difference in CNV length using a custom Python script.

Inversions were taken as the consensus between Delly and Lumpy with the same restrictions and filtering steps implemented for CNVs. Translocations were called under similar approaches as CNVs and inversions. The differences are no length minimums, position differences between Lumpy and Delly calls being a maximum of 1,000 base pairs, and MAPQ scores of 50 or greater are included for analysis. To reduce false positives, these translocation calls were then merged with gene break calls made by CNVKIT by gene name. CNVKIT gene breaks were called using the “breaks” option under default options.

Every previously discussed somatic variant calling option included the FVB_NJ WT reference as the normal sample.

### Pathway Analysis

Previously published gene expression data from MMTV-Myc and MMTV-Neu mouse model tumors were downloaded from the GEO DataSets under the accession number GSE15904. The gct converted file containing gene symbols as row names was used as input for ssGSEA available as a module in GenePattern^82^. Pathway analysis was conducted using the MSigDB gene set databases c1 (positional), c2 (curated), and c6 (oncogenic signature) with default settings. The resulting output data matrix with pathway enrichment scores for each sample was employed for downstream hierarchical clustering and graphical representations.

### Mutation Verification

Mutations observed in the whole genome sequencing analysis for APC, RARA, and KIT genes were screened and confirmed by Sanger sequencing for matched tumor samples and other tumors for a total of 15 samples (5 EMT, 5 squamous, and 5 microacinar tumors). Briefly, RNA was extracted from mammary tumors using the Qiagen RNeasy Midi KIT. cDNA was generated using the AppliedBiosystems High-Capacity cDNA Reverse Transcription KIT. Genes were amplified under PCR using the following primers: c-KIT 5’ ATAGACTCCAGCGTCTTCCG 3’ and 5’ GCTCCCAATGTCTTTCCAAAACT 3’; RARA 5’ TTGTGCATCTGAGTCCGG TT 3’ and 5’ TGGGCAAGTACACTACGAACA 3’; APC 5’ CCGCTCGTATTCAGCAGGT 3’ and 5’ CCTGCAGCCTATTCTGTGCT 3’. PCR parameters are standard parameters as listed on New England BioLabs PCR protocol (M0273) V1. PCR fragments were purified using QIAquick PCR Purification KIT or QIAquick Gel Extraction KIT. Sanger sequencing was performed by GENEWIZ/Azenta under standard premixed conditions, with one of the PCR primers used as the sequencing primer. Sequence alignment and visualization were performed with Geneious Prime-2020.0.5 software.

### Circos Plots

Representative mouse circos plots for each histological subtype were generated using Circos^83^ as a software package available on MSU’s high performance computing center (HPCC). All mutations, CNVs, inversions, and translocations were mapped at exact mm10 genomic coordinates of each event.

Working from the outermost region of each plot, it begins with a labelled mouse ideogram with chromosomes presented in ascending order. The next innermost rings consist of SNVs color coded to represent low (yellow), moderate (orange), or high (red) predicted impact as determined by Metect2. Following, the next inner ring depicts copy number variations as whole integer copy number changes, with height corresponding to copy number integers of one or two. Copy number gains are depicted in red, while copy number losses are depicted in blue. In the final innermost circle, translocations are colored randomly to one of the two chromosomes involved in the translocation event. Inversions are colored in black. Only somatic variants that satisfy the previous requirements for somatic variant calling are included for each tumor.

### Copy Number and Gene Expression Correlation

Correlation of copy number and gene expression was done on CNVKIT. After generating copy ratios and copy segments under the “batch” WGS method, discrete copy number segments were generated using the “segment” method. The “cnv_expression_correlate” method was used to generate Kendall rank correlation coefficients and Pearson correlation coefficients on discrete copy number data for all 9 tumors that underwent WGS and gene expression data for all 42 tumors that underwent transcriptomic profiling. Correlation coefficients for each gene were then mapped onto Circos plots, with significant Kendall’s τ coefficients (≥ 0.3) colored blue and in the outermost ring. In the innermost ring, significant Pearson’s r coefficients (≥ 0.7) are colored red. Non-significant values for both metrics were colored black. Correlation coefficients could only be generated for genes with discrete copy number changes at +/-1 of ploidy level or greater differences, so a majority of the genome will show no correlation.

### Unsupervised Clustering Analsyses

All unsupervised hierarchical clustering was performed using the clustermap function as part of the Seaborn Python library. Distance between clusters was computed using the Ward variance minimization algorithm.

### PCA

Principal component analysis was performed using the scikit-learn^84^ library, utilizing the StandardScaler and PCA packages. The number of principal components analyzed was 2 in every instance. Results were visualized using a custom scatter plot in matplotlib.

### Copy Number Heatmaps

All heatmaps displaying log2 fold change of copy number segmentation data were generated using CNVKIT, both for initial copy number segmentation and copy number ratio file generation, as well as visualization.

### Mutational Signatures and Mutational Burden

Mutational burden plots were generated in matplotlib. Mutational signatures were derived using the deconstructSigs^41^ R package and plotted using matplotlib.

WGS data for MMTV-Neu and MMTV-PyMT previously analyzed in the lab and used in mutation plots are available under the NIH SRA with BioProject number PRJNA541842. Processing of Neu and PyMT WGS fastq reads was done according to the same methods as the MMTV-Myc samples.

### Volcano Plot

The volcano plot of ssGSEA c6 oncogenic signatures for the EMT subtype compared to both squamous and microacinar signatures was plotted in Python using BioinfoKIT^85^. The log-fold change threshold is at 0.4 and the p-value threshold is set at 0.05.

### Kaplan Meier Curves

Kaplan Meier plots were made with using the Survminer R package. Publicly available and deidentified TCGA nonredundant breast cancer patient data was used to construct Kaplan Meier curves using criteria stated for each figure.

### t-SNE Plot

The t-SNE diagram was created using the t-SNE implementation inside of scikit-learn. t-SNE was performed on the optimized 32 gene subset expression data of the PAM50 gene set as determined by RFECV using a support vector machine classifier. The resulting scatterplot of data is generated using matplotlib and colored according to the legend.

### Supervised Machine Learning

The supervised machine learning soft voting classifier implemented using scikit-learn consists of logistic regression, support vector machine, random forest, XGBoost, and multi-layer perceptron classifier probabilities merged together. METABRIC gene expression data for the 32 genes with matched PAM50 subtypes was shuffled and split 70-30 into training and test data sets, respectively. To combat overfitting and training on biased data, PAM50 subtype proportions were kept intact and probability predictions averaged over 15 instantiations.

### Other Visualizations

Bar chart visualizations and pairplots were generated using Matplotlib, Seaborn, and Yellowbrick.

### Human Data Usage

Clustering was performed on human breast cancer transcriptional data matched with intrinsic subtypes from the anonymized METABRIC dataset, which is readily available through cBioPortal. Kaplan Meier analyses were performed on non-redundant human breast cancer patients from all breast cancer studies available in cBioPortal as of time of publication that match the criteria specified in each plot.

### Statistical Considerations

Unless otherwise stated, statistical tests with p-value displayed are done using a two-sided student’s t-test.

### Software Versions

BioinfoKIT-2.1.0, Bokeh-2.4.3, BWA-mem-0.7.17, Circos-0.69.6, CNVKIT-0.9.9, Delly-0.7.8, FASTQC-0.11.7, GATK-4.1.4.1, Lumpy-0.2.13, Matplotlib-3.4.3, Mutect2-2.1, NumPy-1.20.3, Pandas-1.3.4, Panel-0.13.1, Picard-2.18.1, Python-3.9.7, SAMtools-1.9, scikit-learn-0.24.2, SciPy-1.9.0, Seaborn-0.11.2, SnpEff-4.3, ssGSEA-10.0.11, Trimmomatic-0.38, VarScan-2.4.1, and Yellowbrick-1.5.

## Declarations

### Ethics Approval and Consent to Participate

No human subjects or clinical specimens were involved in this study. All mouse investigations were approved by the institutional animal care and use committee (IACUC) at MSU.

### Consent for Publication

Not applicable.

### Availability of Data Materials

Mouse MMTV-Myc whole genome sequence data obtained in this study is available under BioProject PRJNA945899. Python and R code used in this study to process data and generate figures is available on GitHub (https://github.com/CarBroke). Other data related to primary and supplementary figure generation are included in supplemental materials.

### Authors’ Contributions

CDB wrote the manuscript, performed whole genome DNA extraction and processing of MMTV-Myc samples, performed all bioinformatic analyses, performed all statistical analyses, performed all machine learning, and contributed to the theoretical design of the study. MMOO performed the mutation verification by Sanger sequencing and ssGSEA processing. MSM contributed to the theoretical and technical design of the machine learning components. ERA performed MMTV-Myc tumor extraction and preservation, performed microarrays on mouse transcriptional data, and lead the theoretical design of the study. All authors contributed to the revision of the manuscript.

## Acknowledgments

The authors thank Jesus Alberto Garcia-Lerena, John Vusich, Marcelio Shammami, Reham Ammar, and James Lord of Michigan State University for insightful discussion.

---

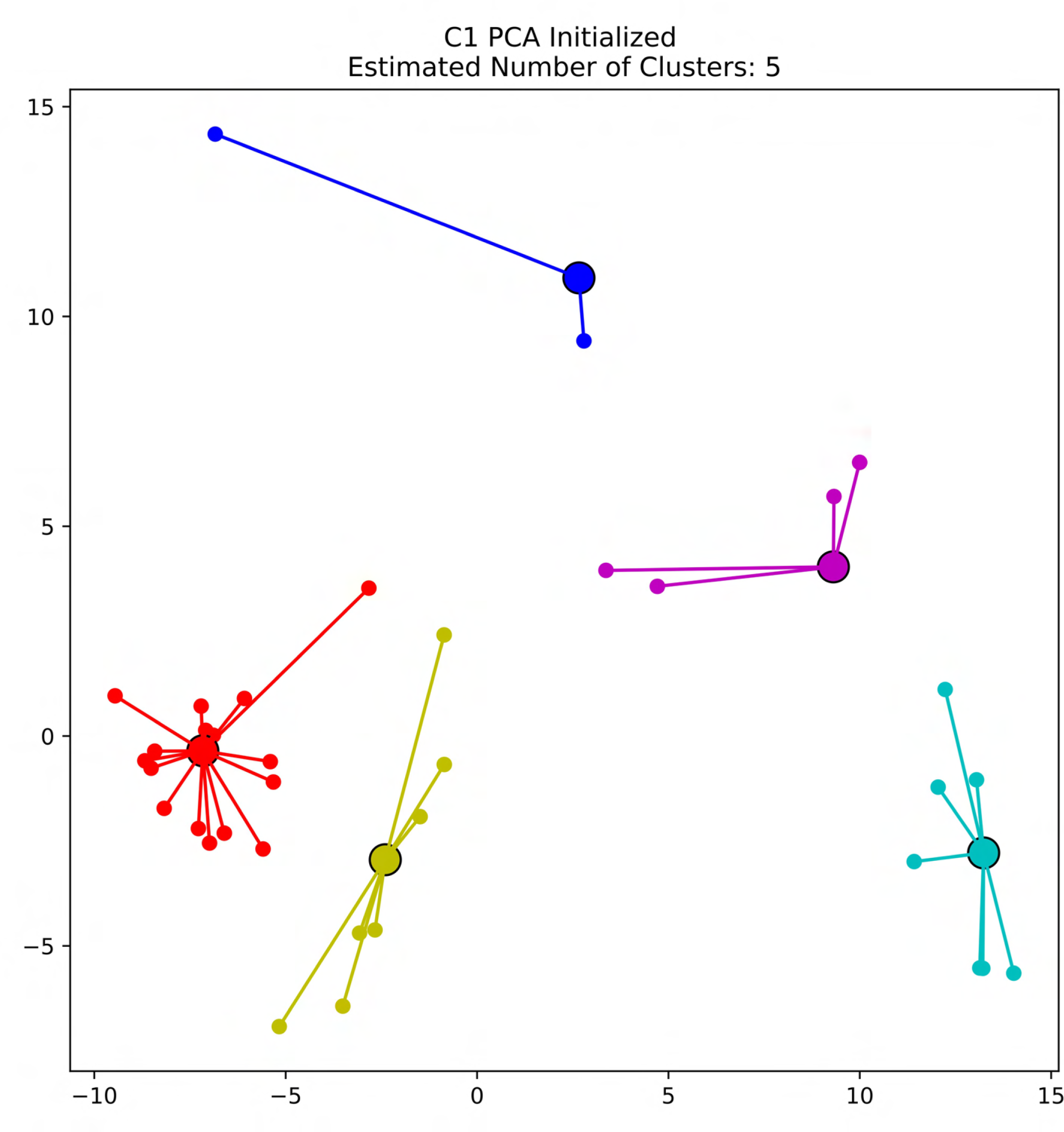

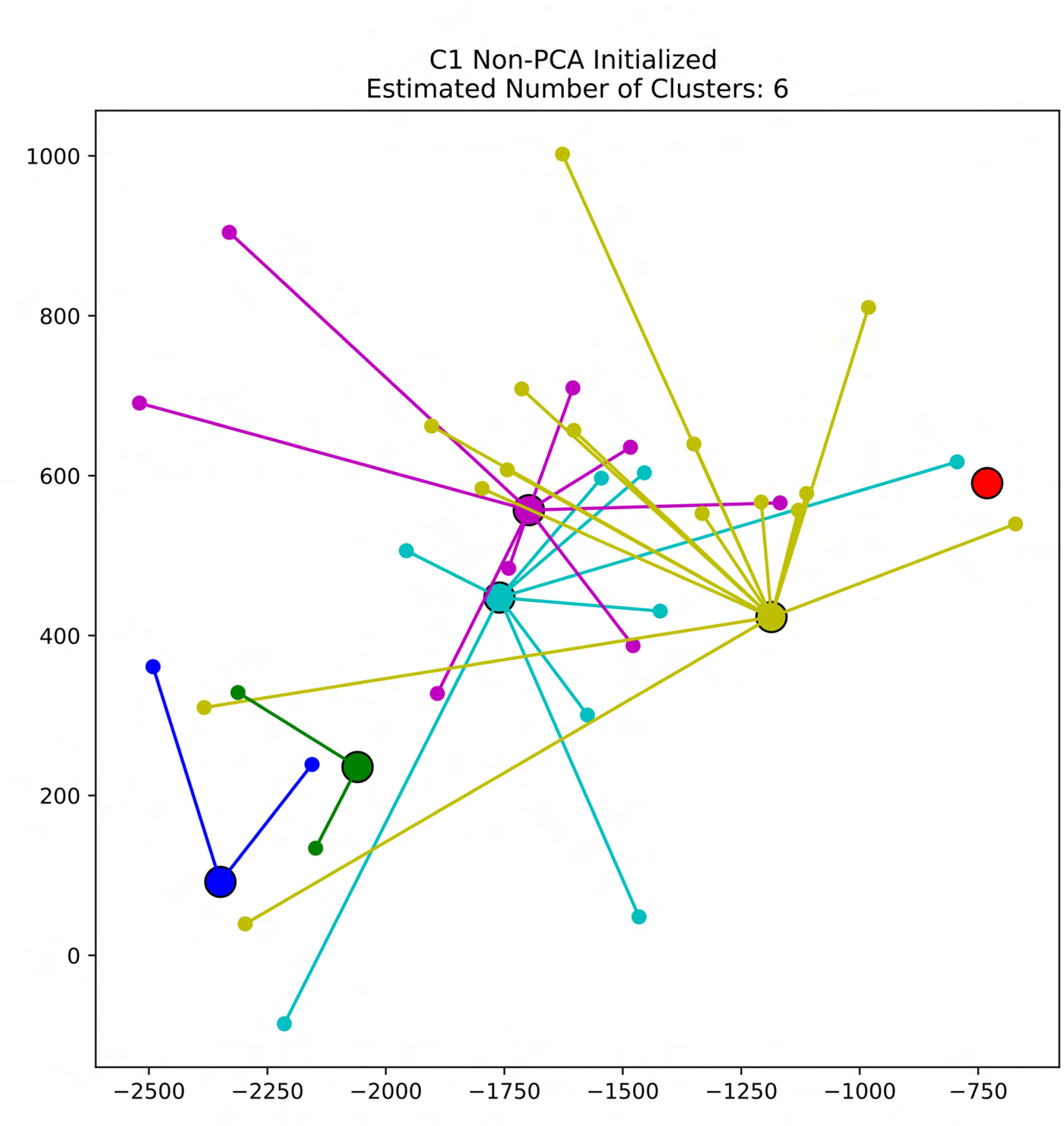

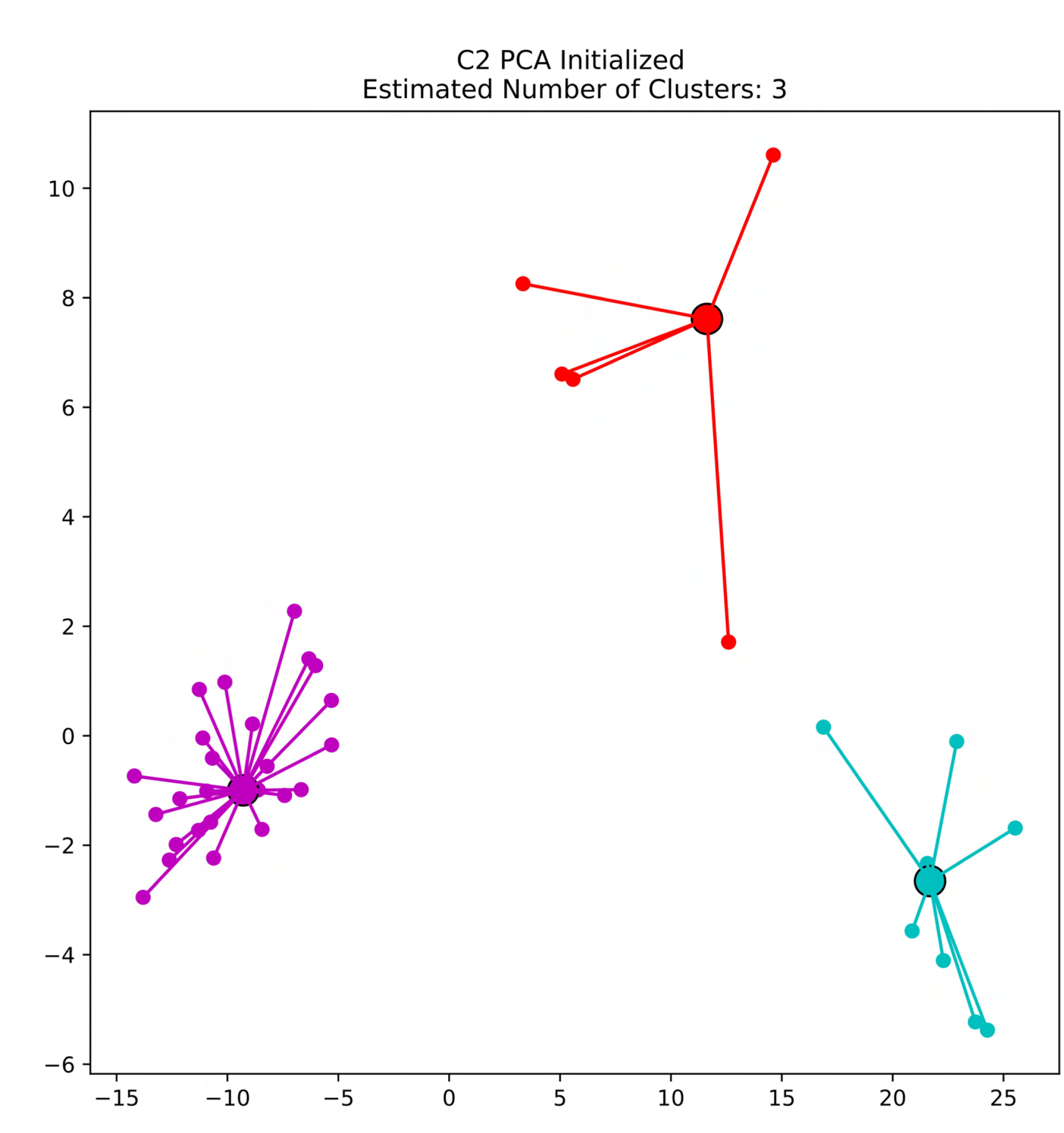

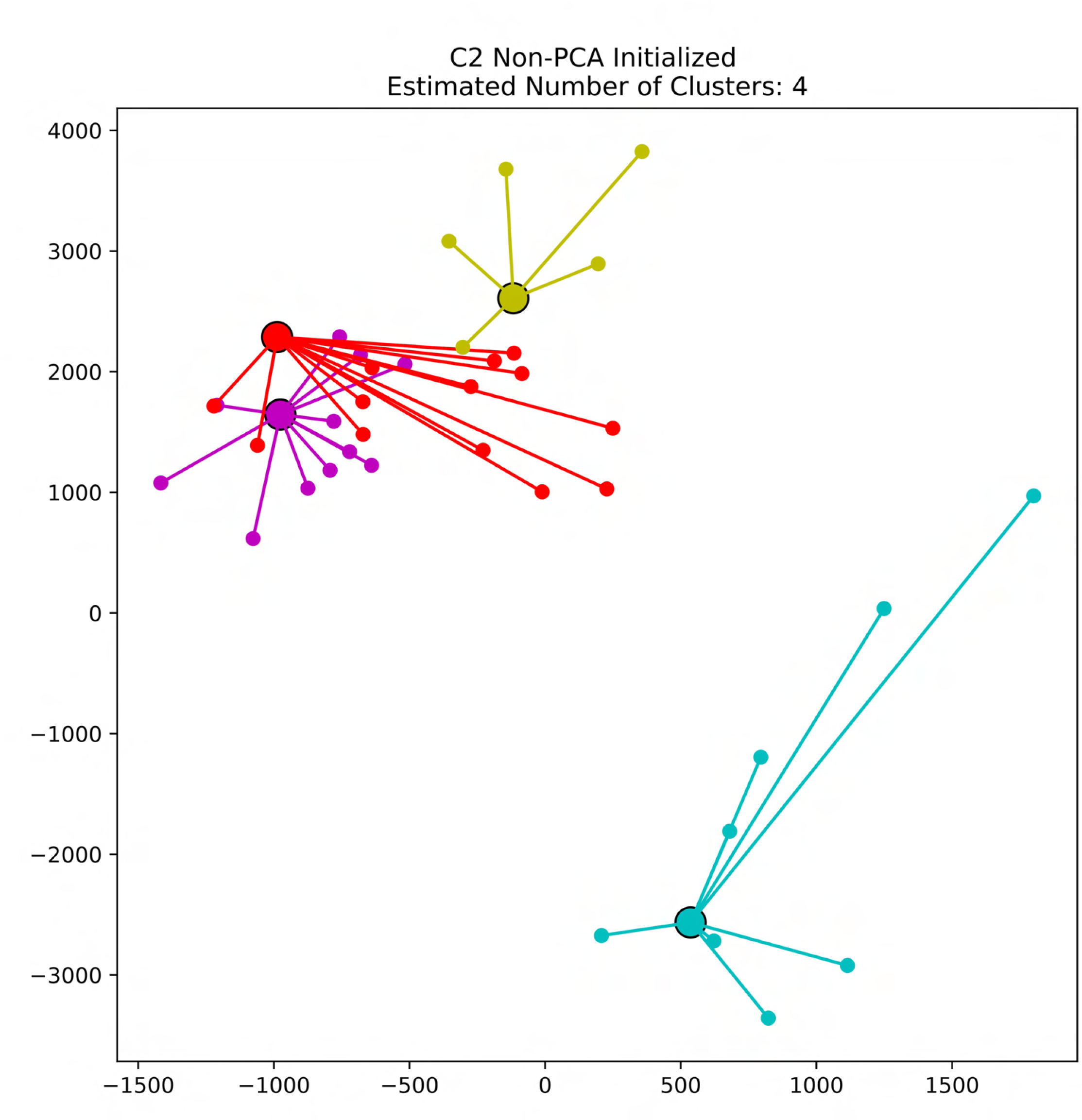

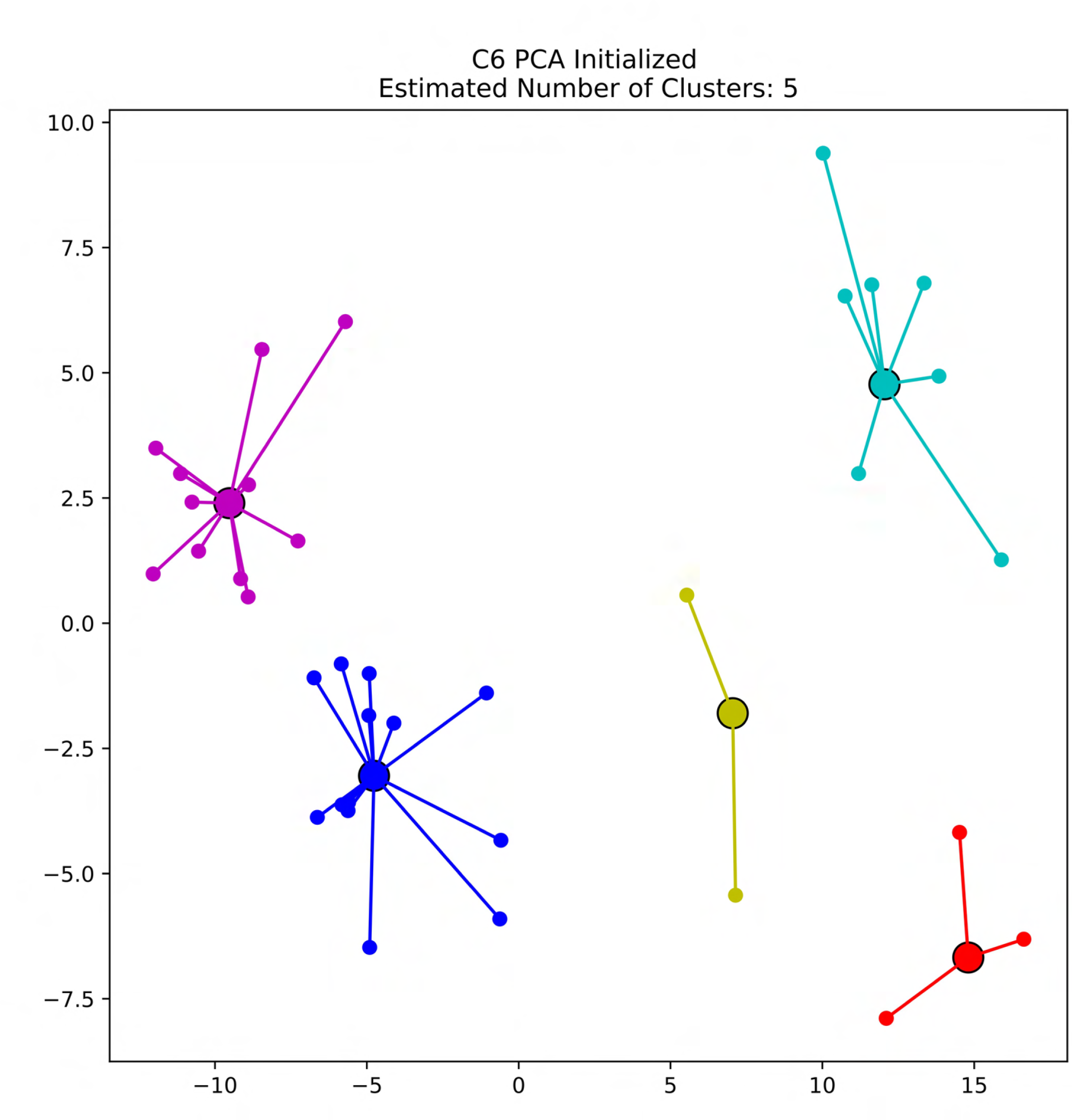

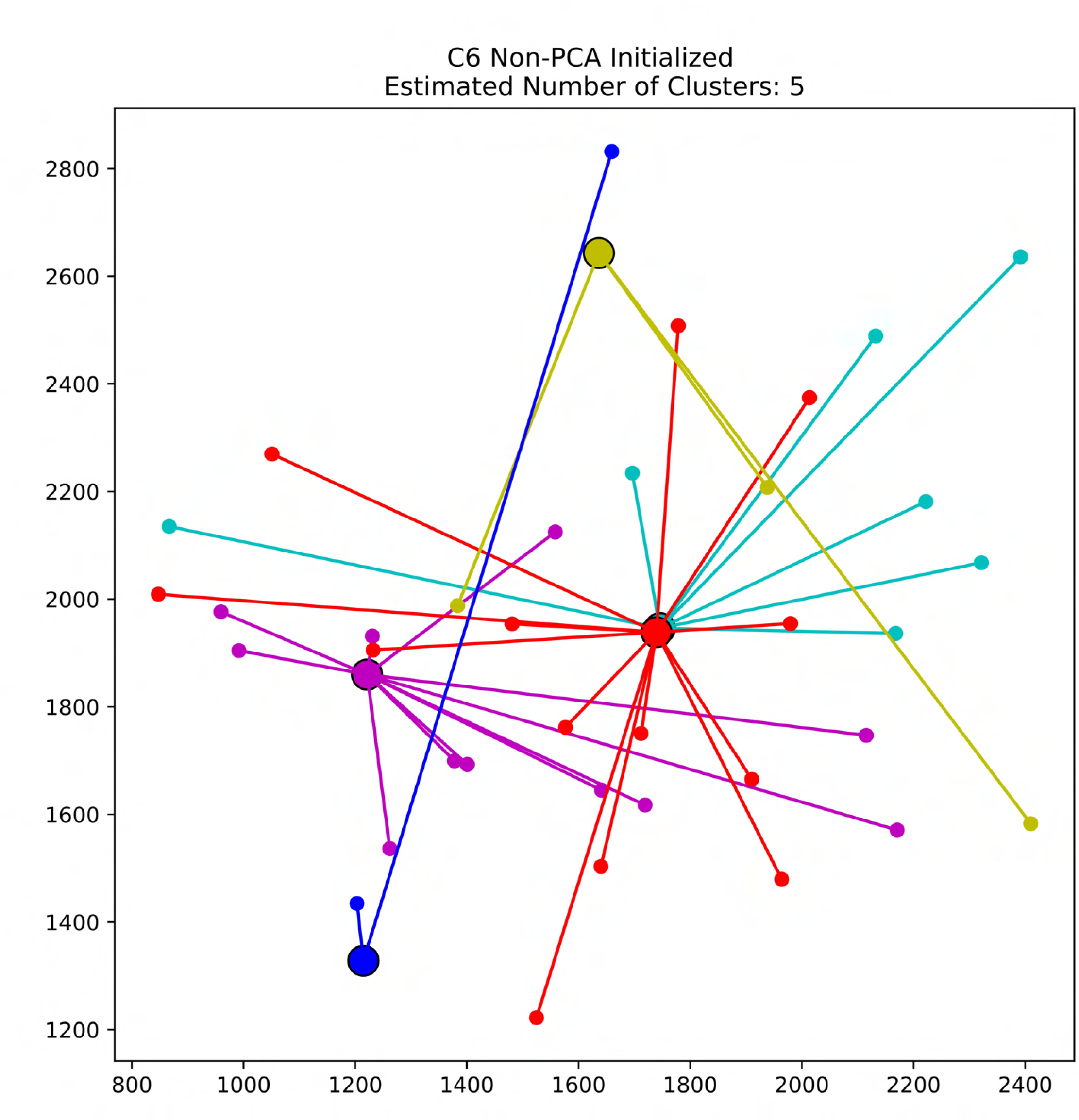

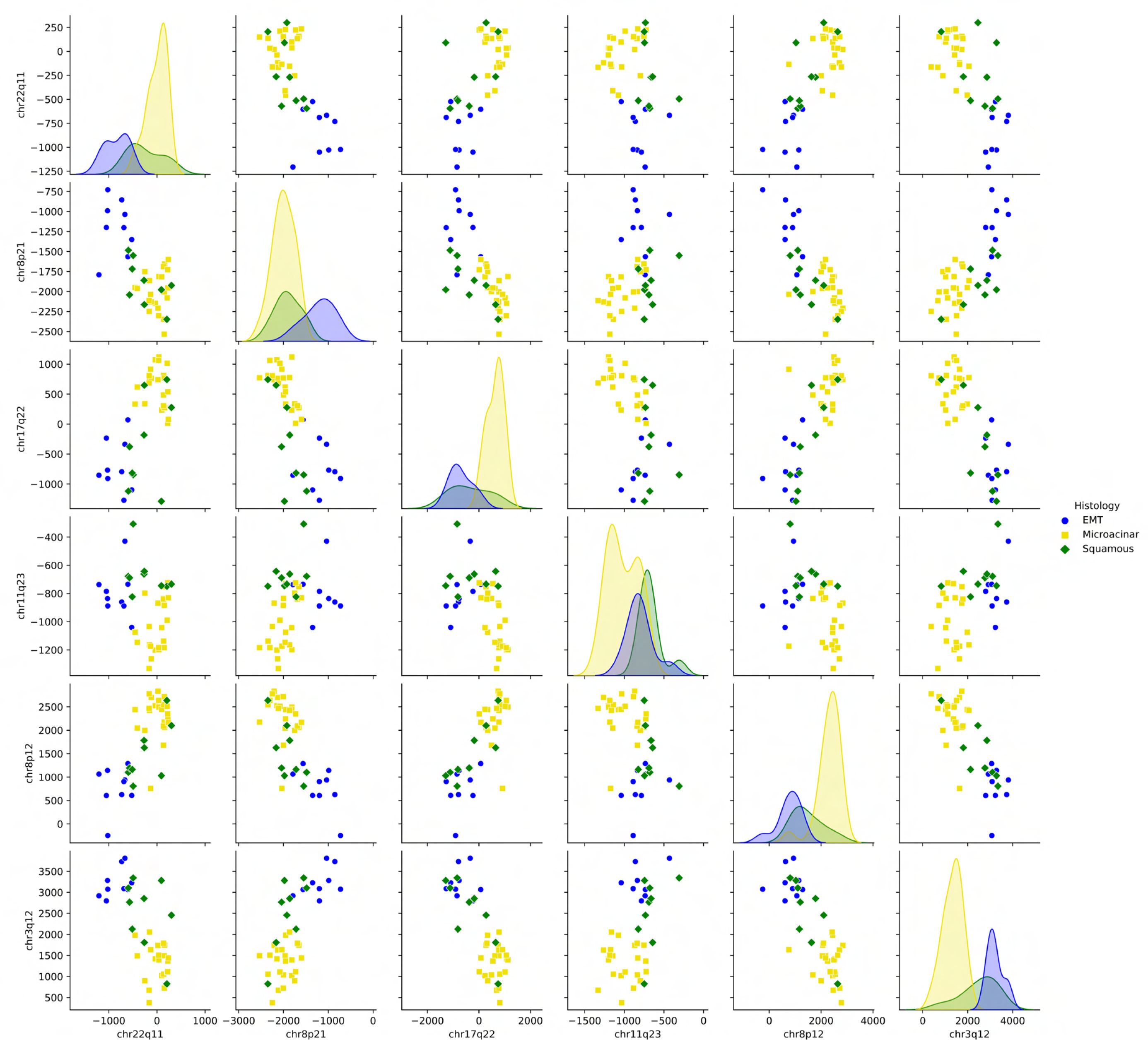

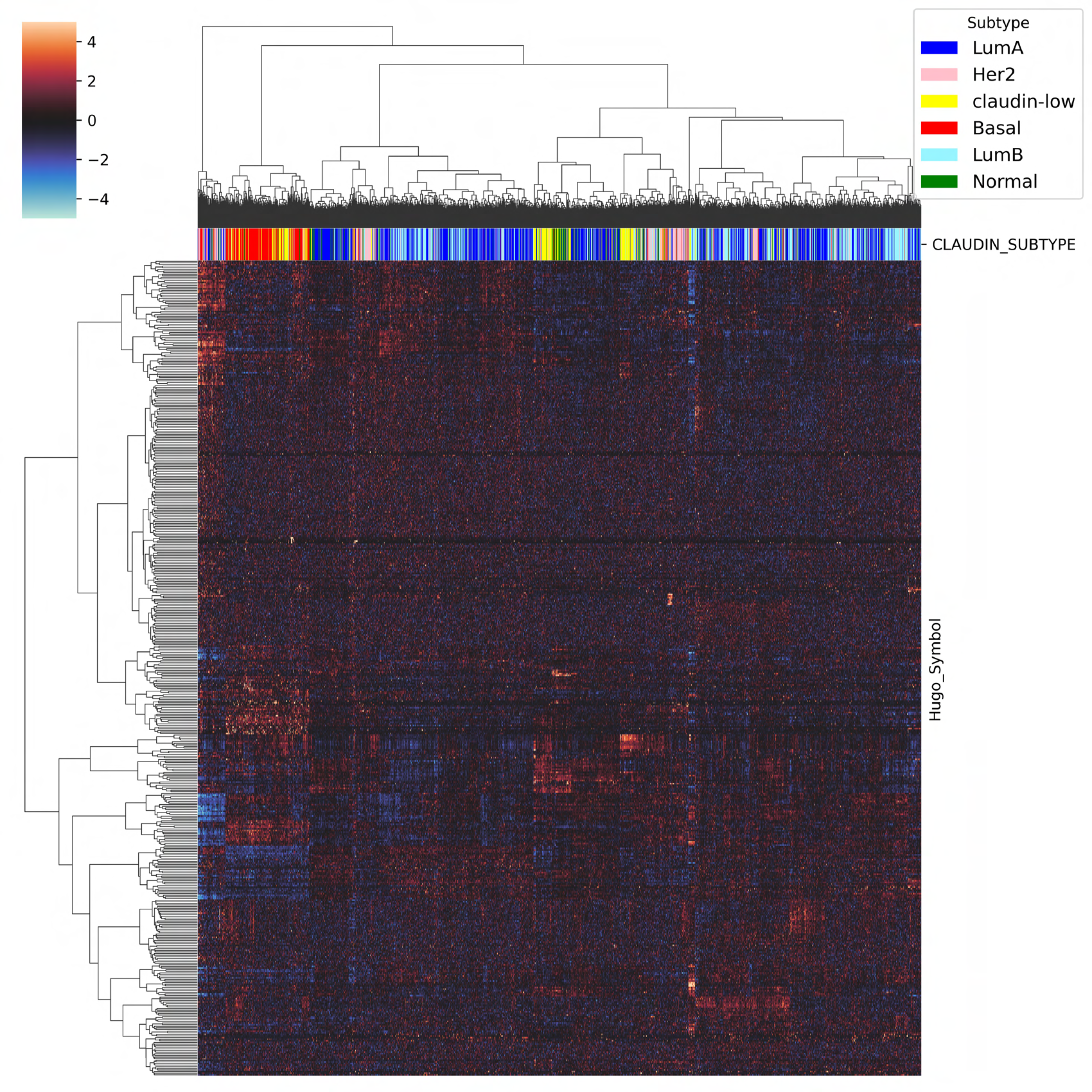

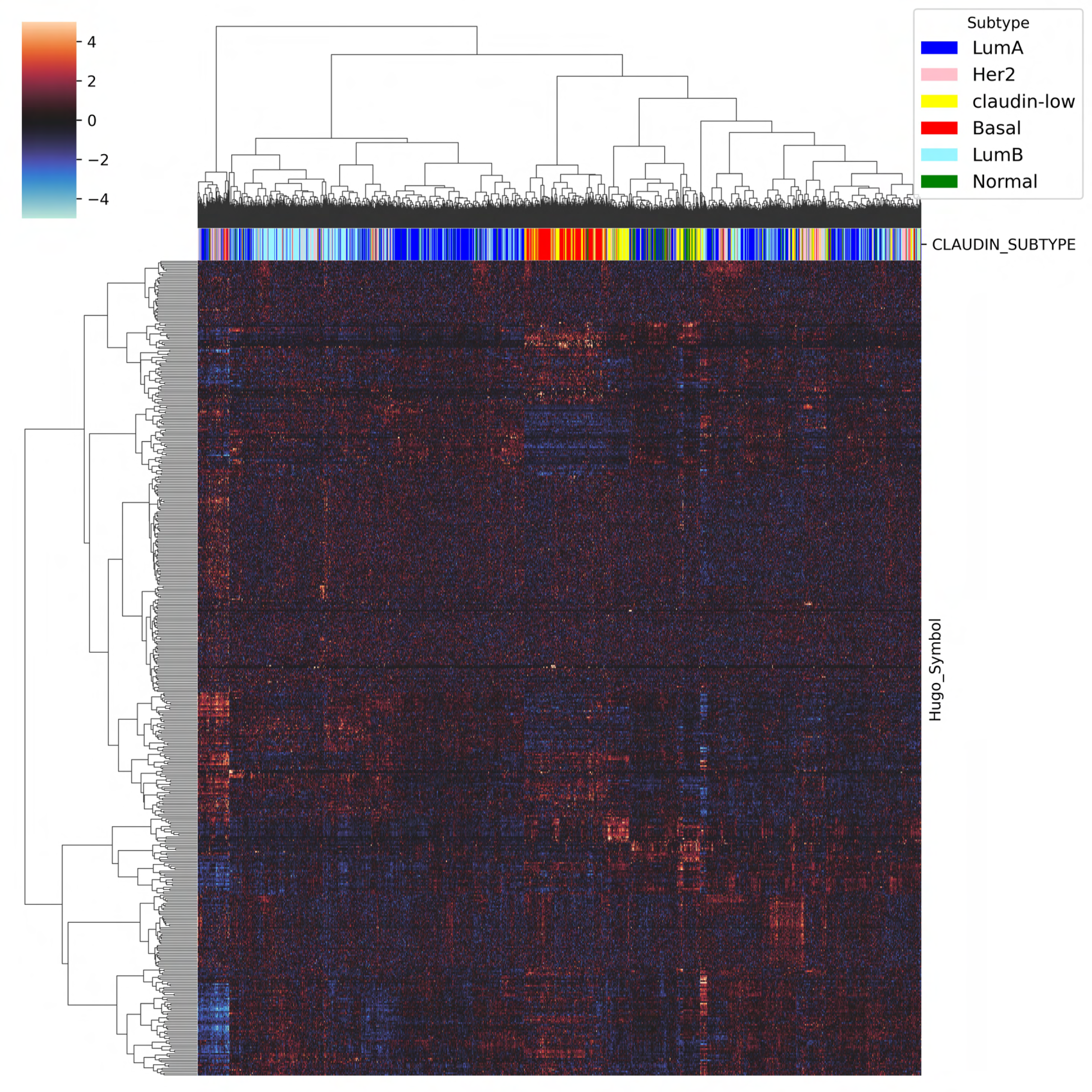

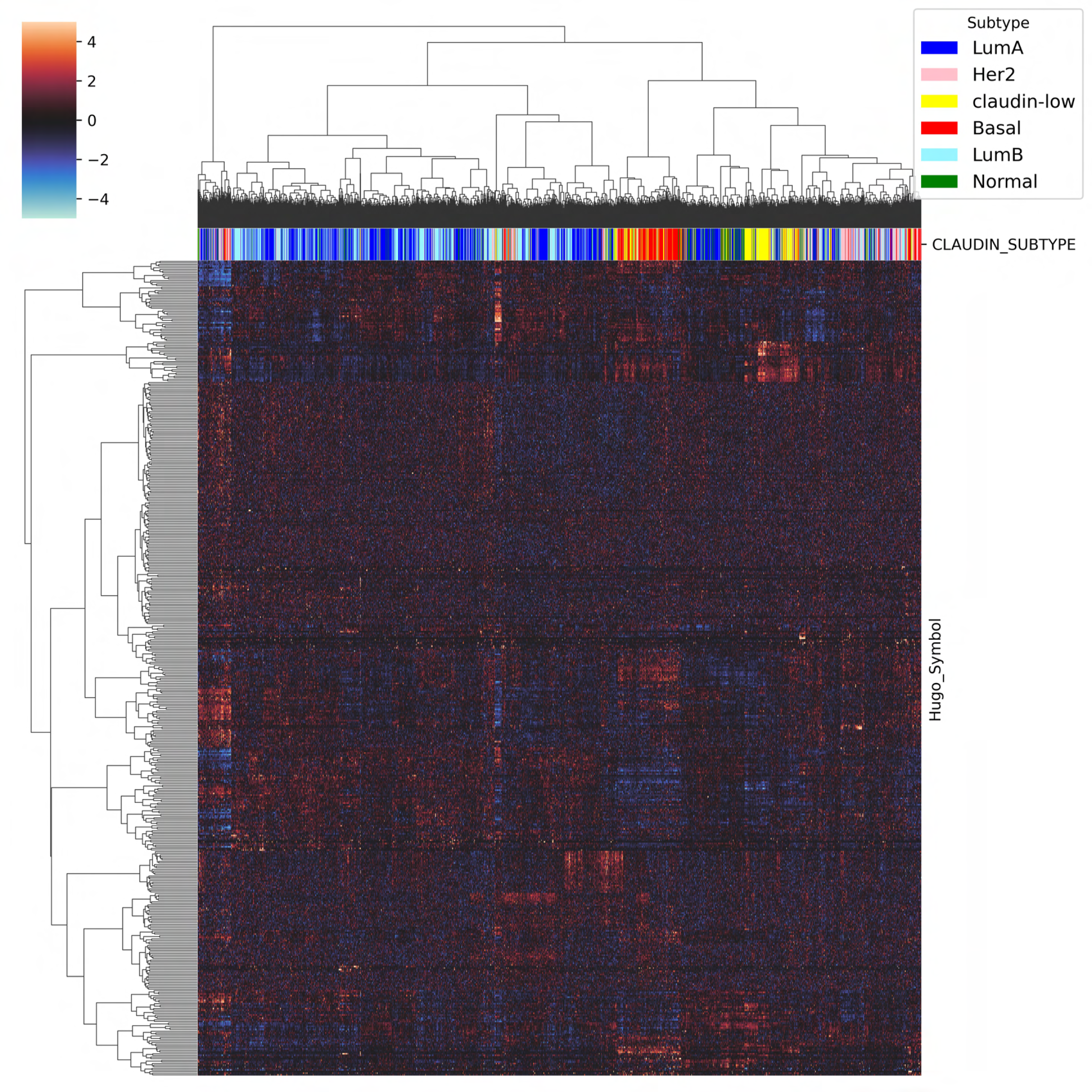

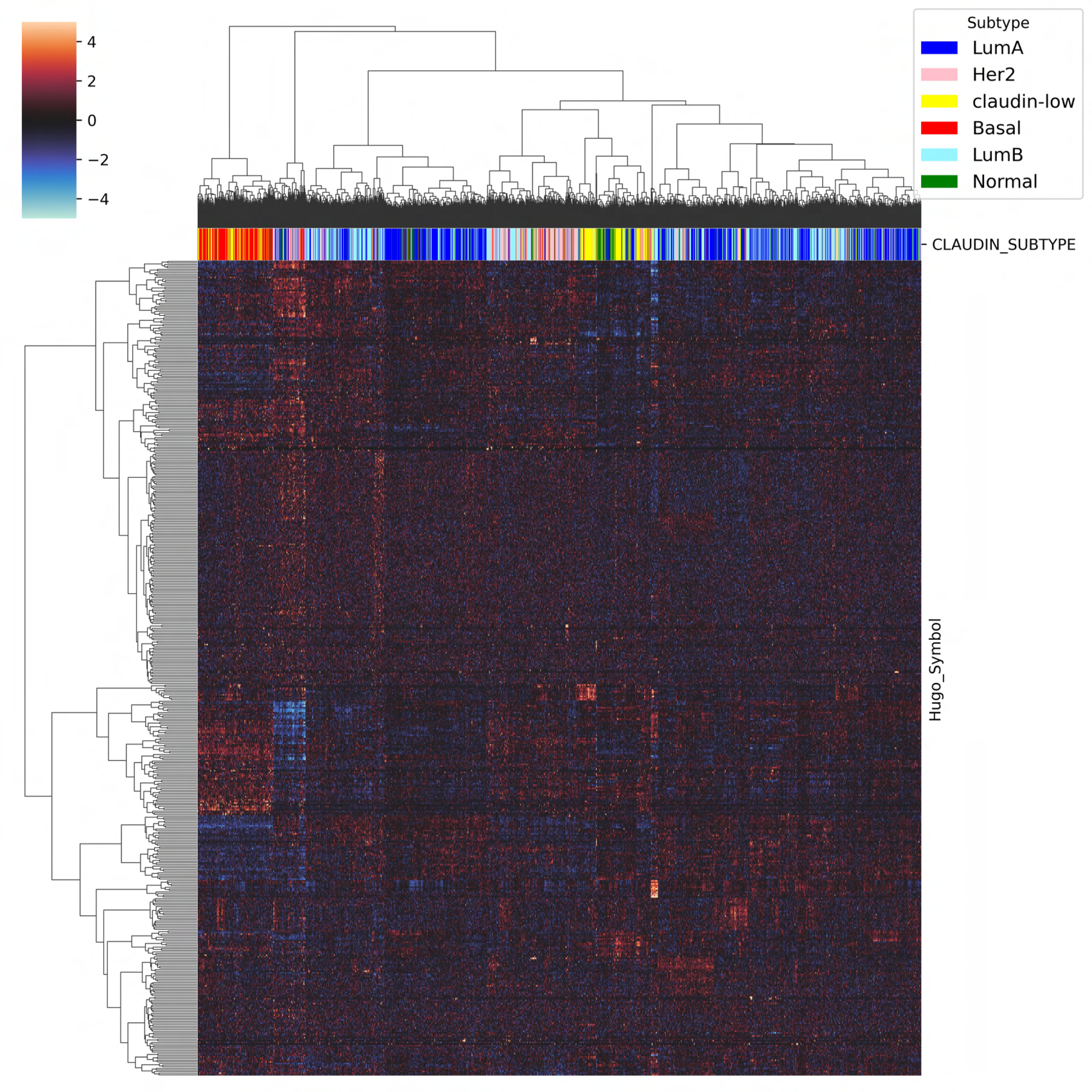

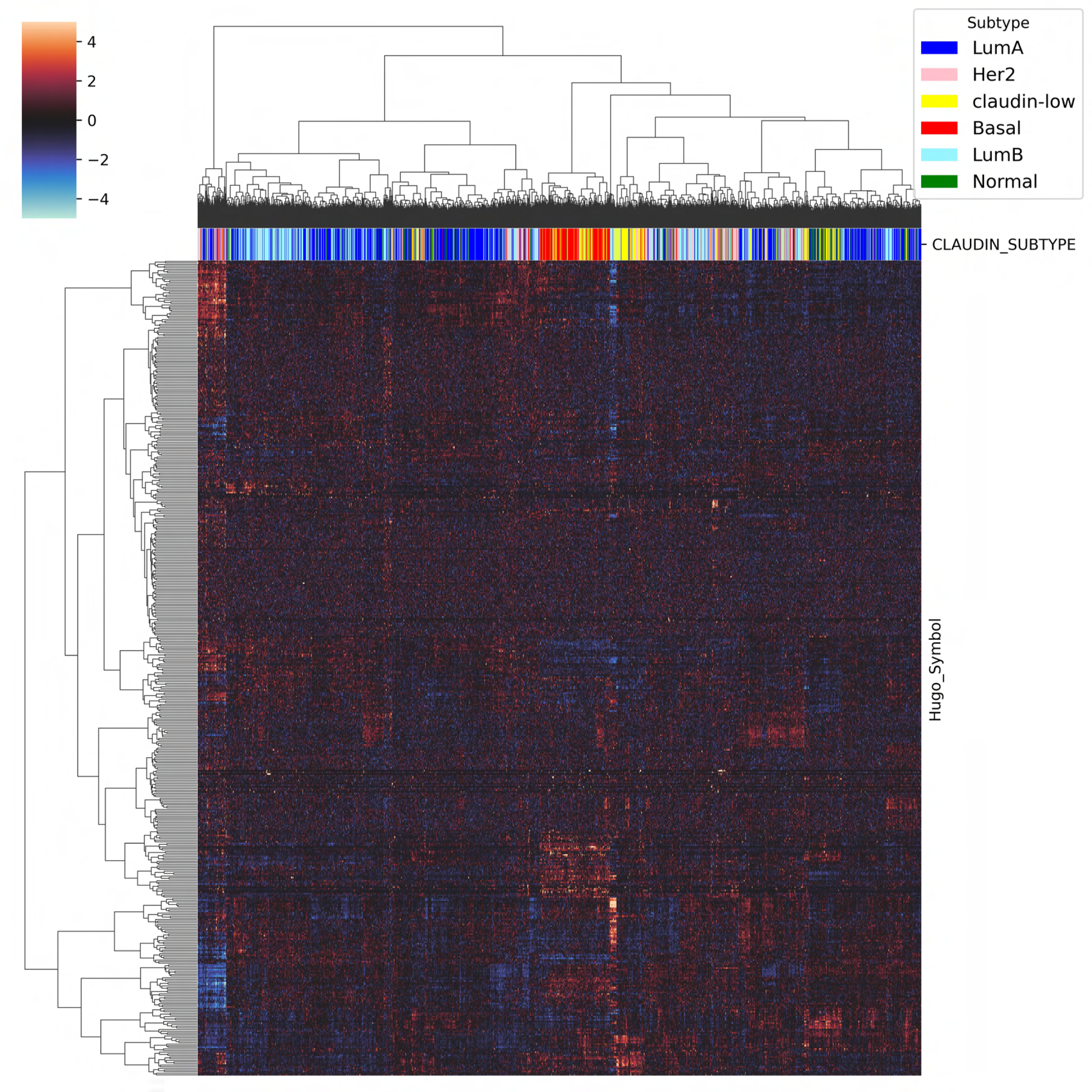

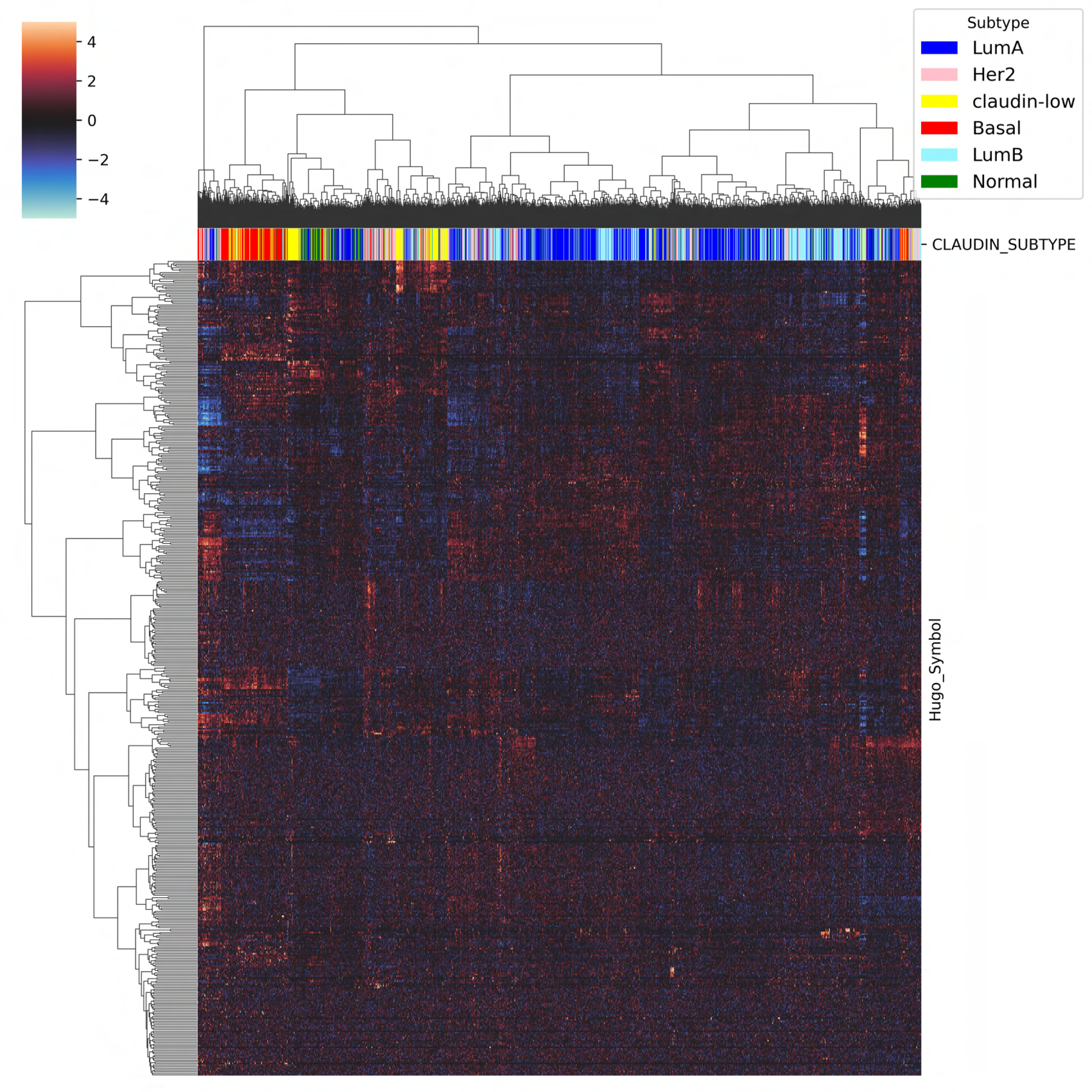

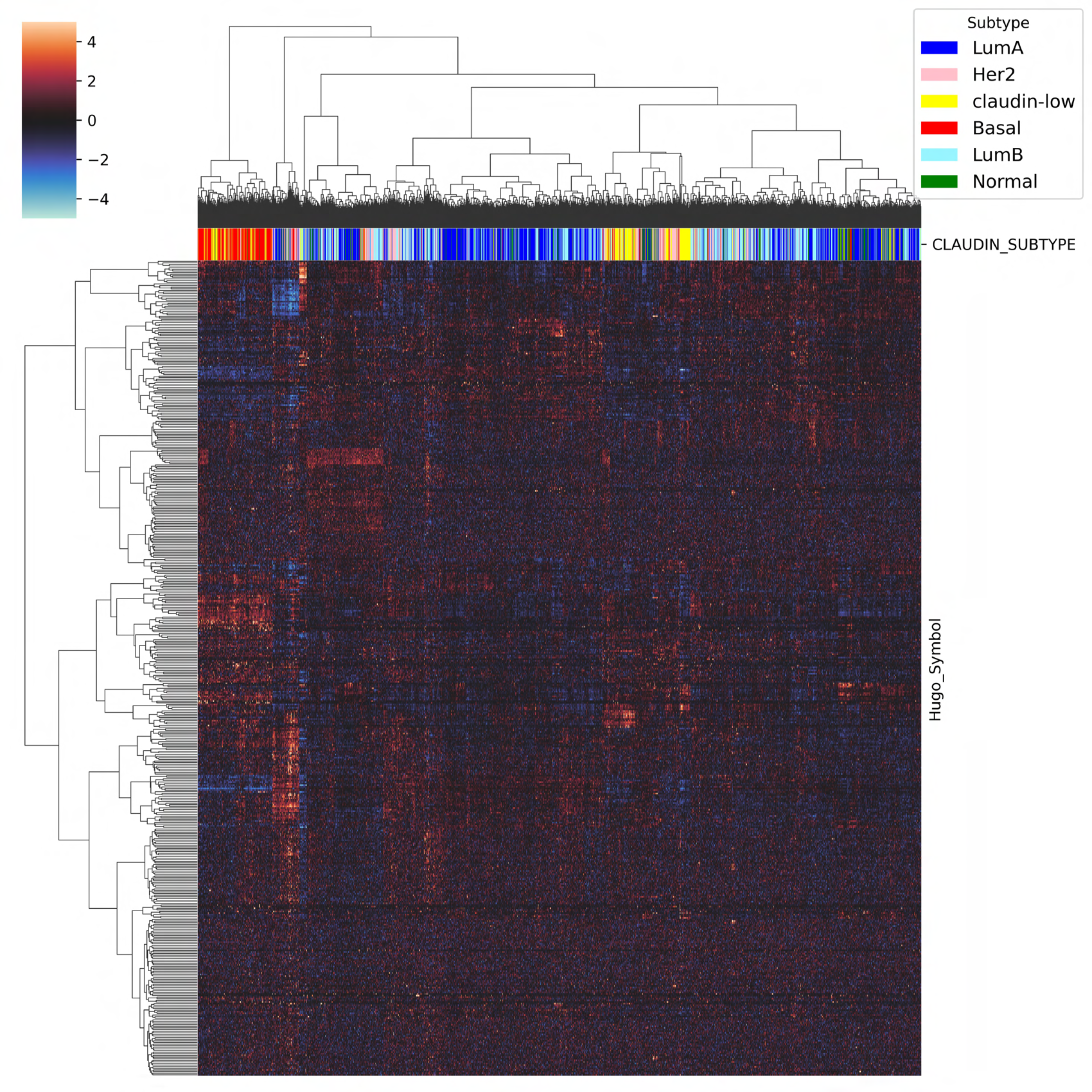

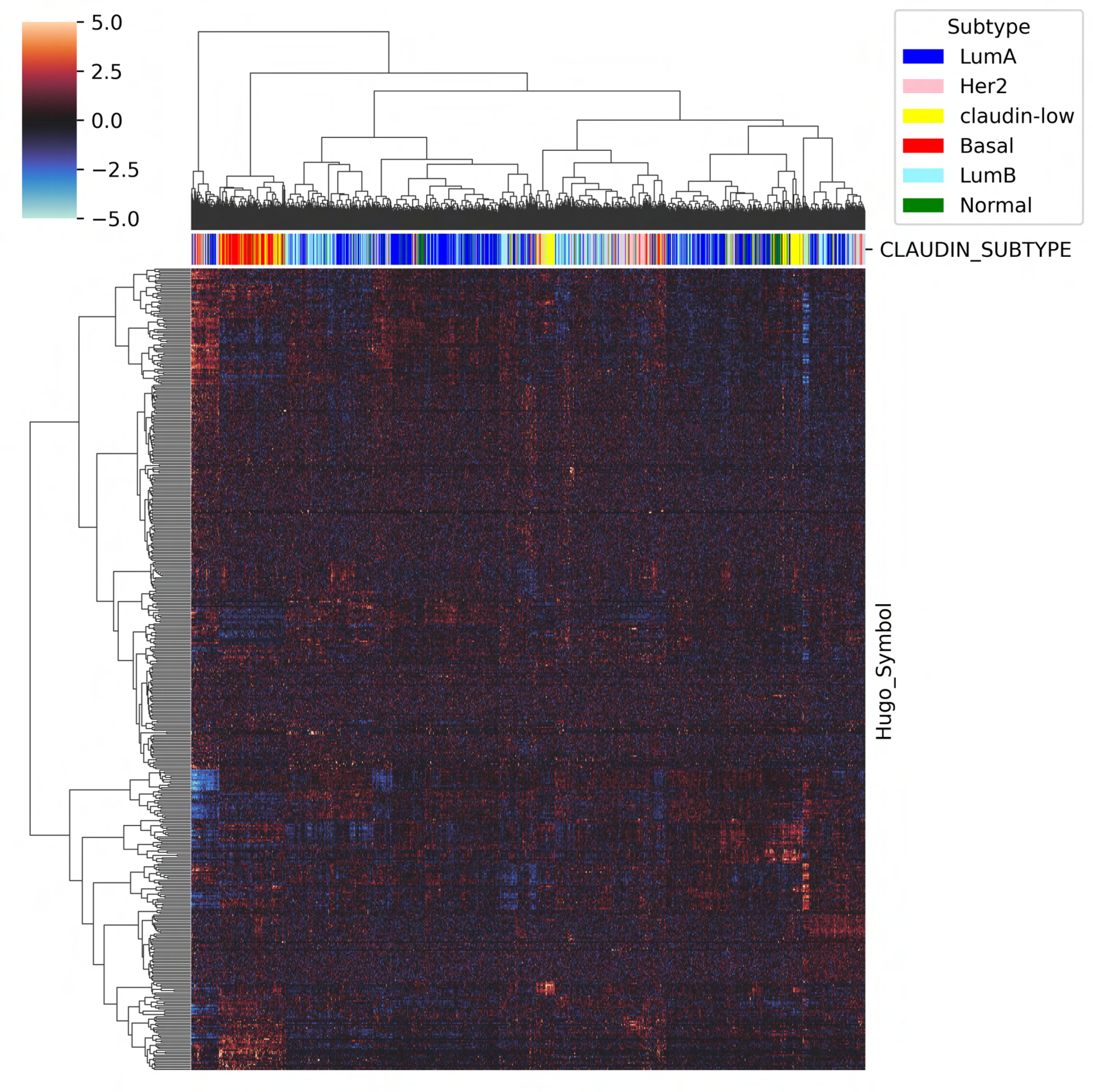

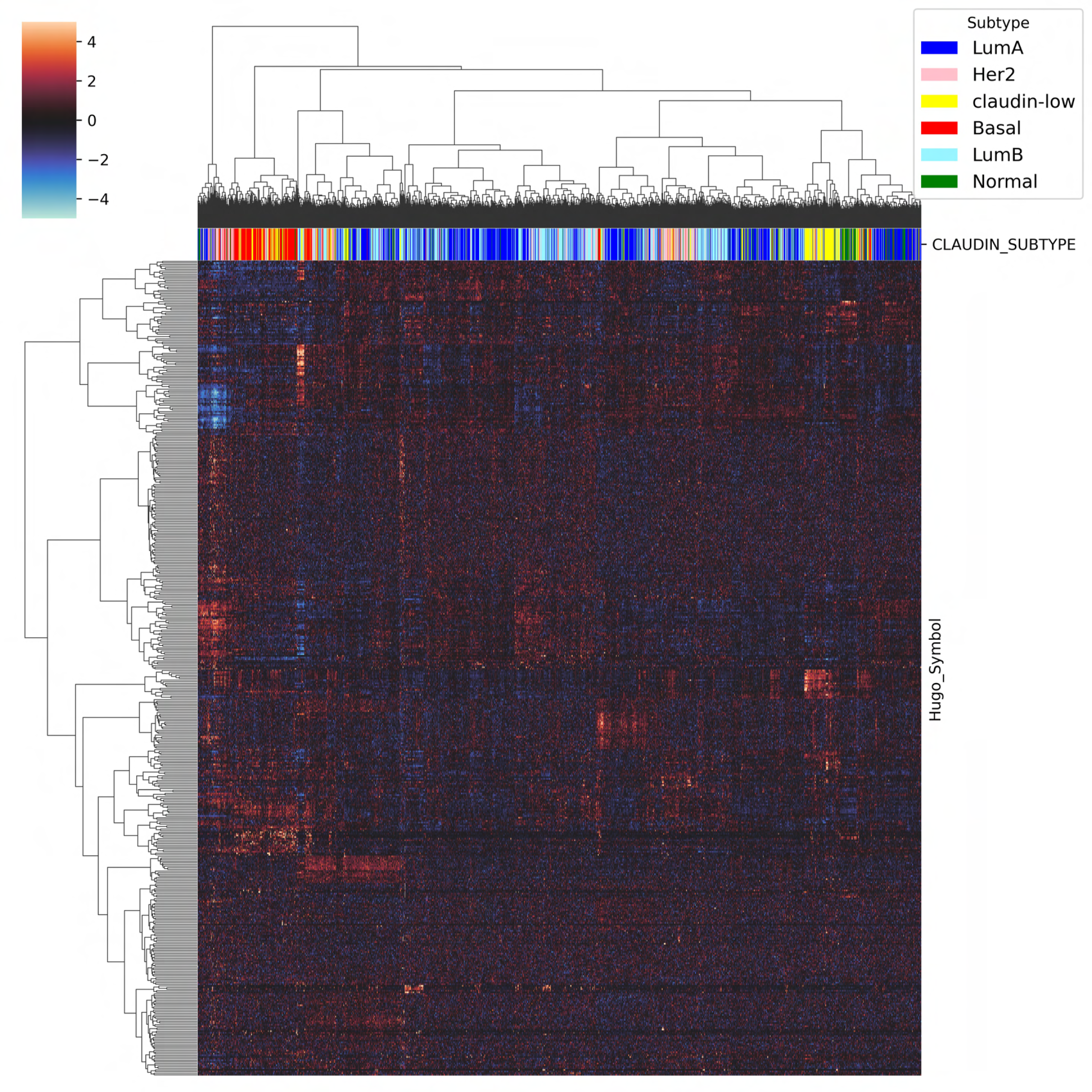

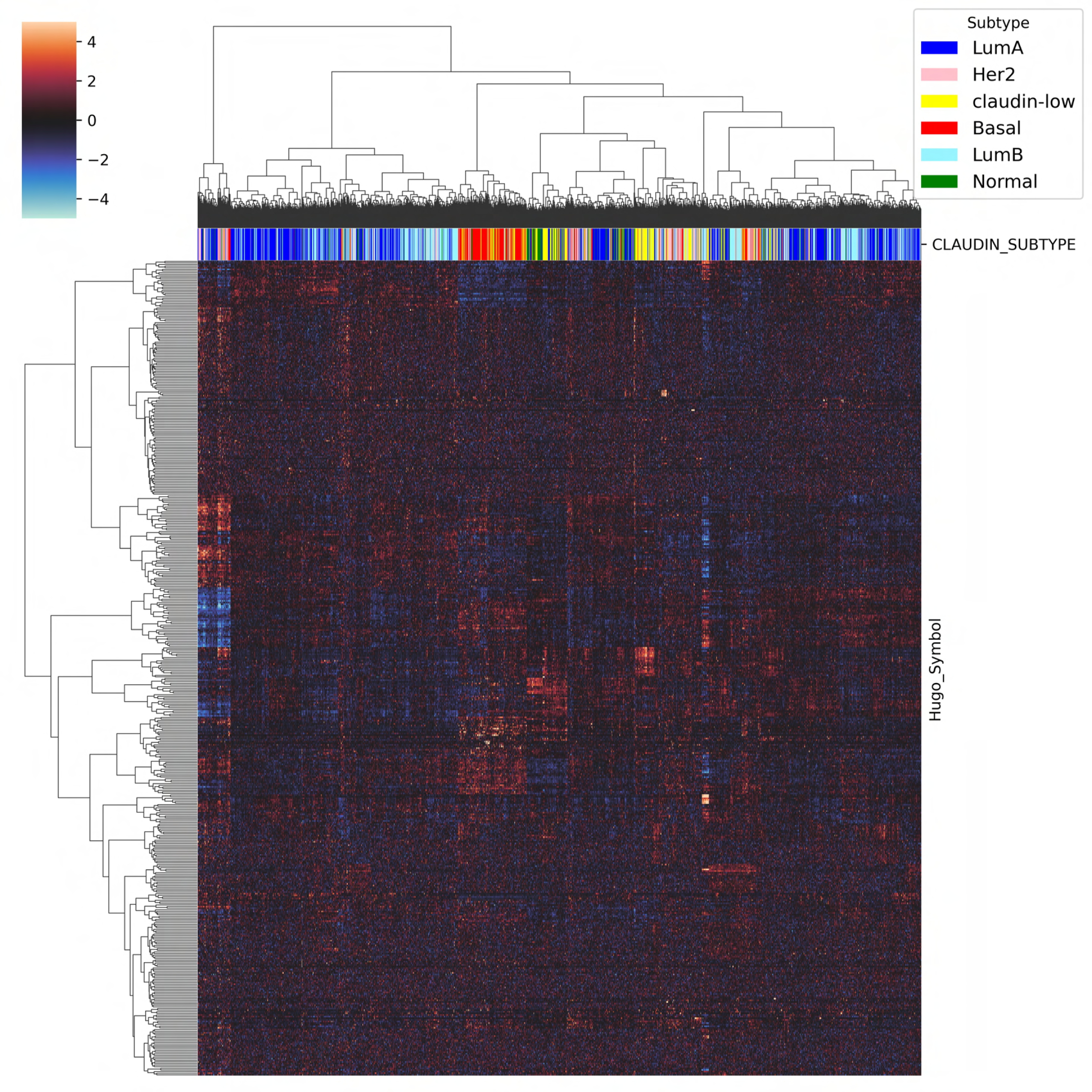

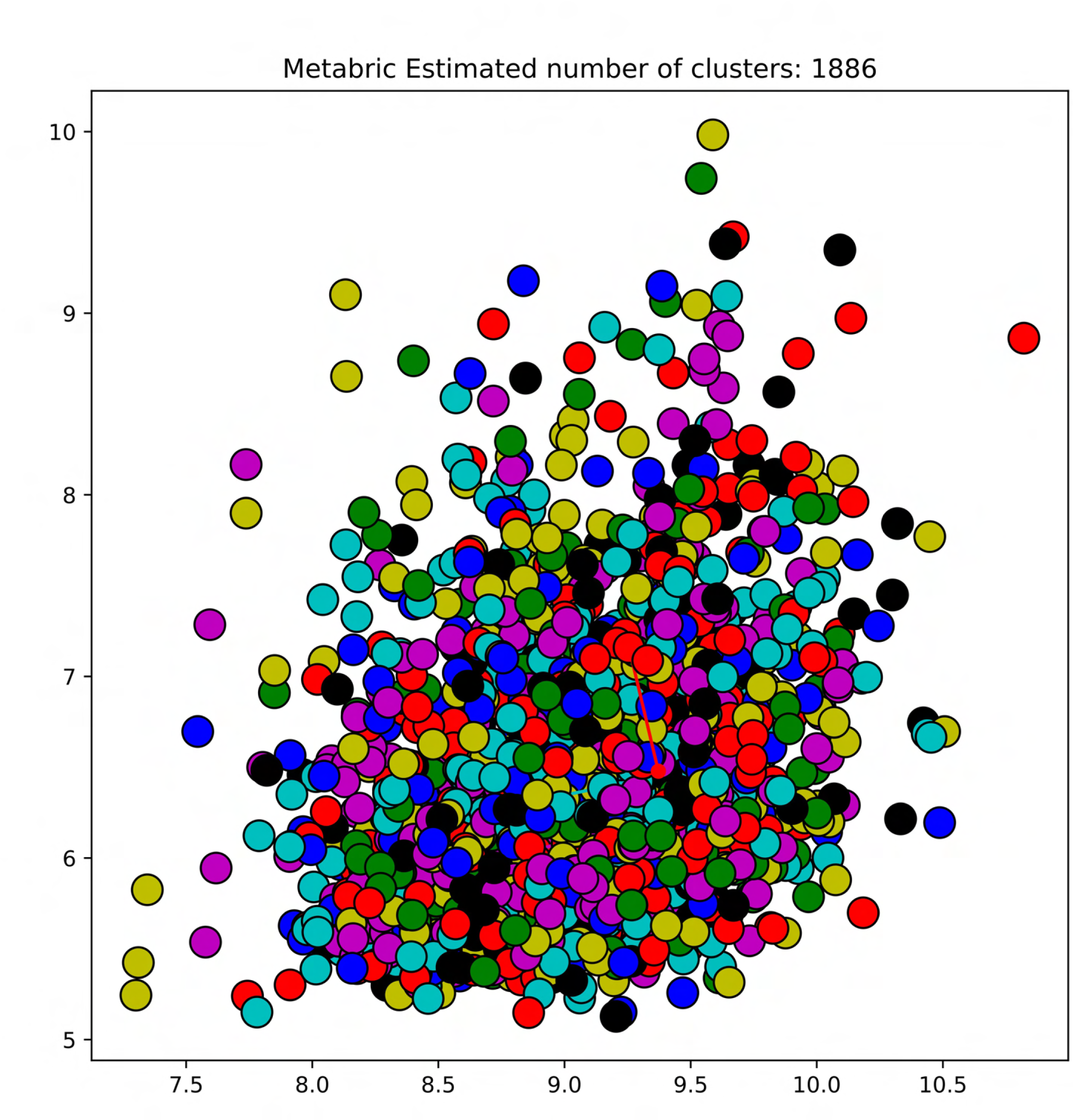

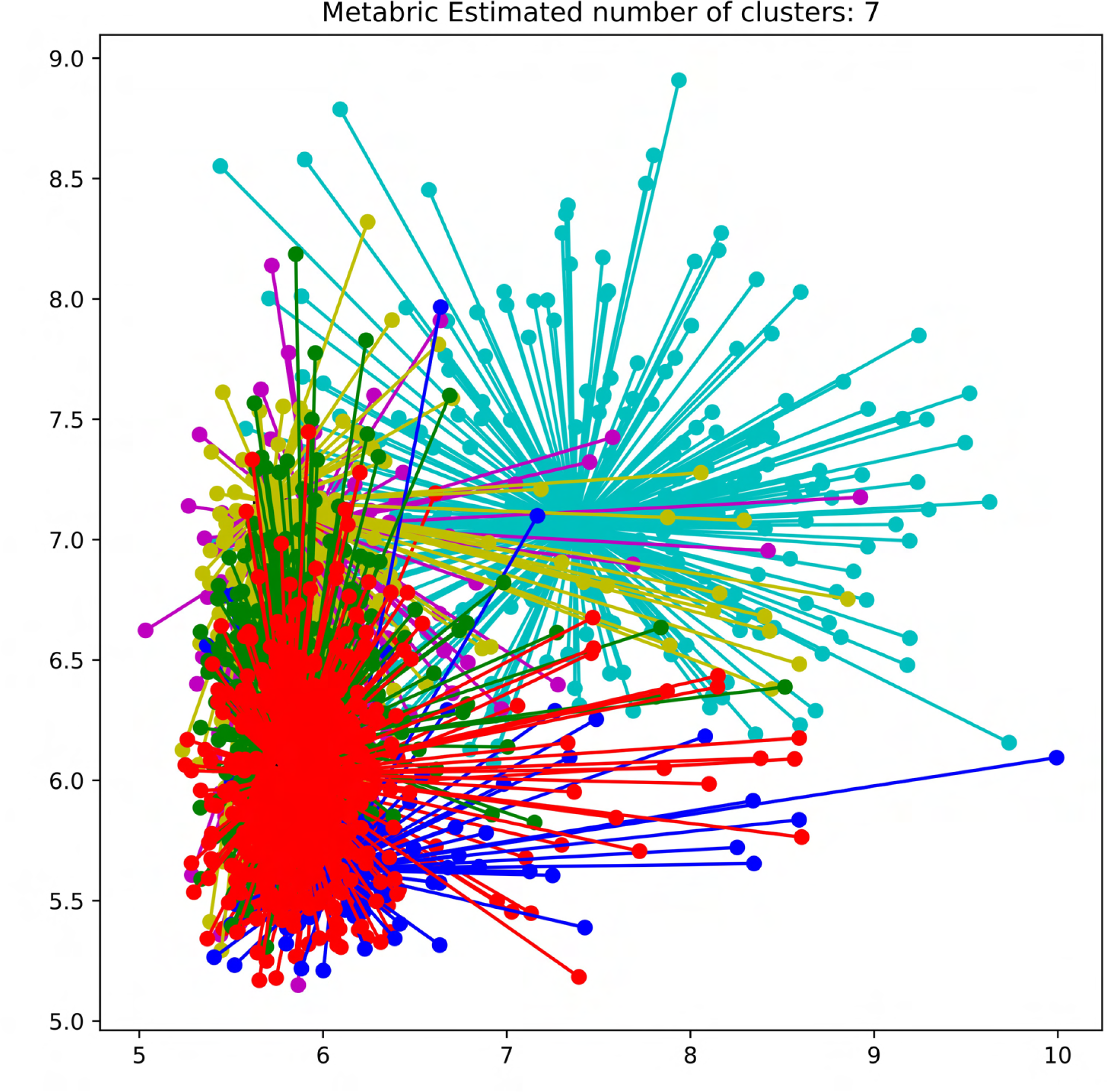

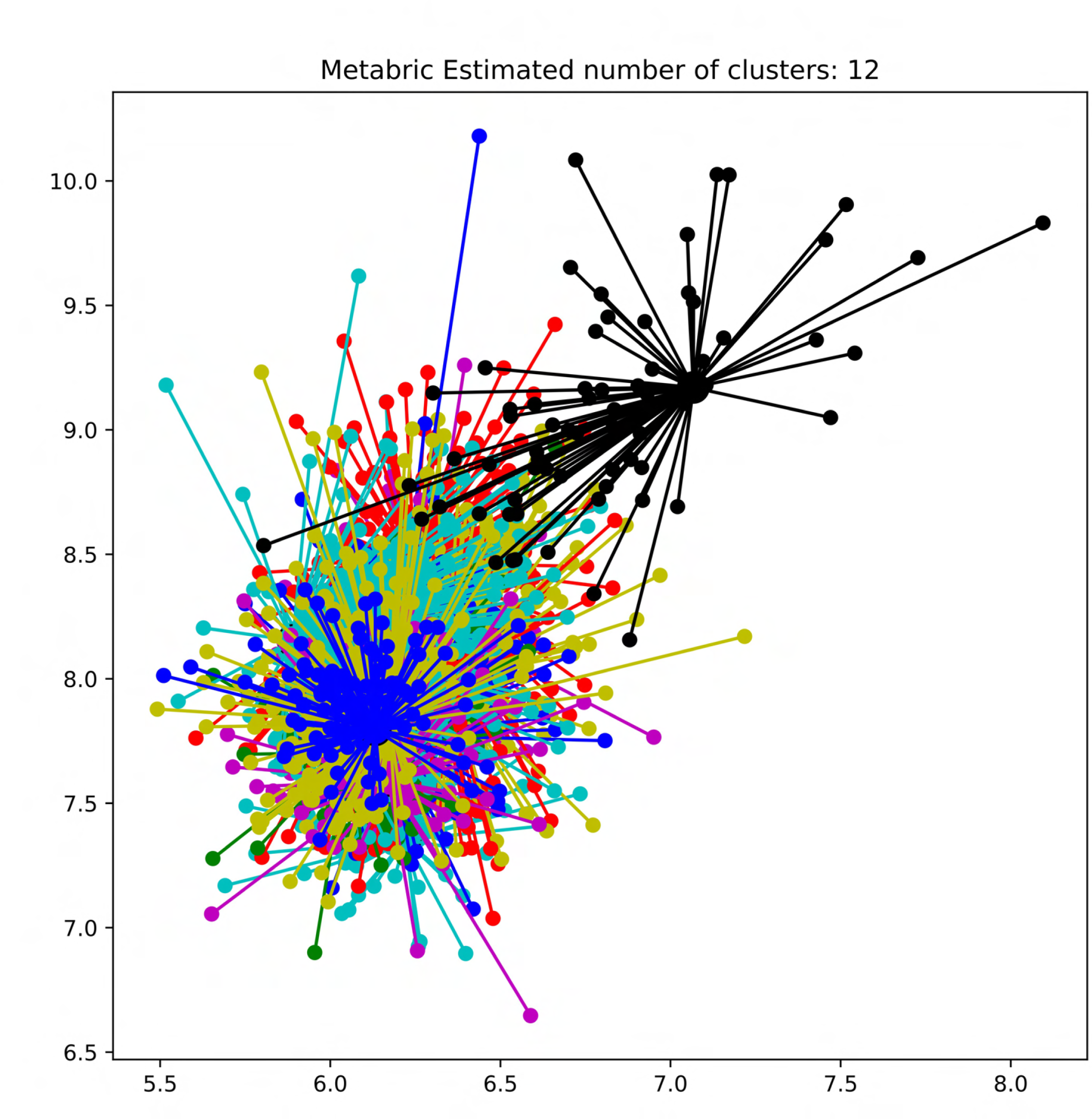

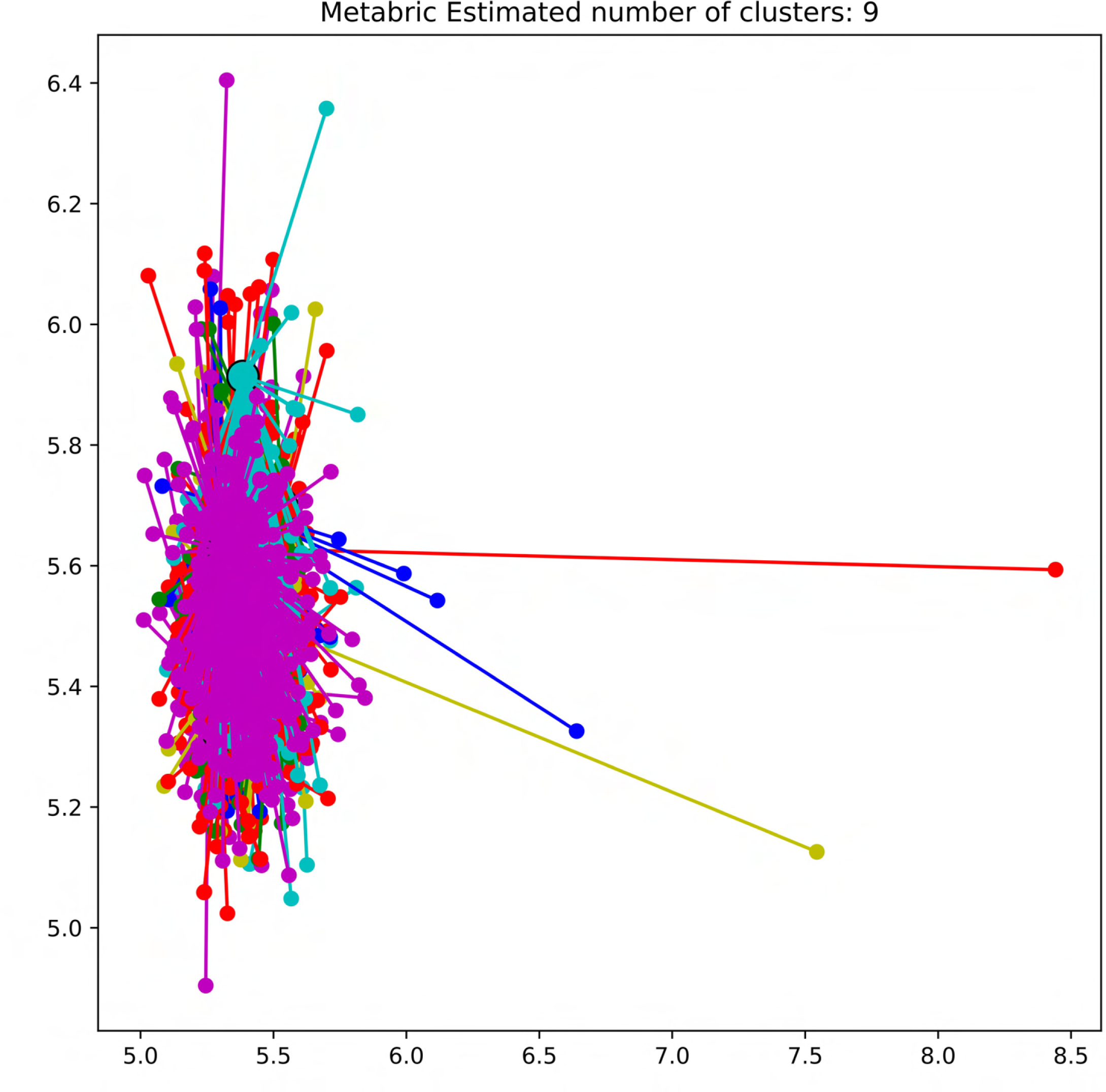

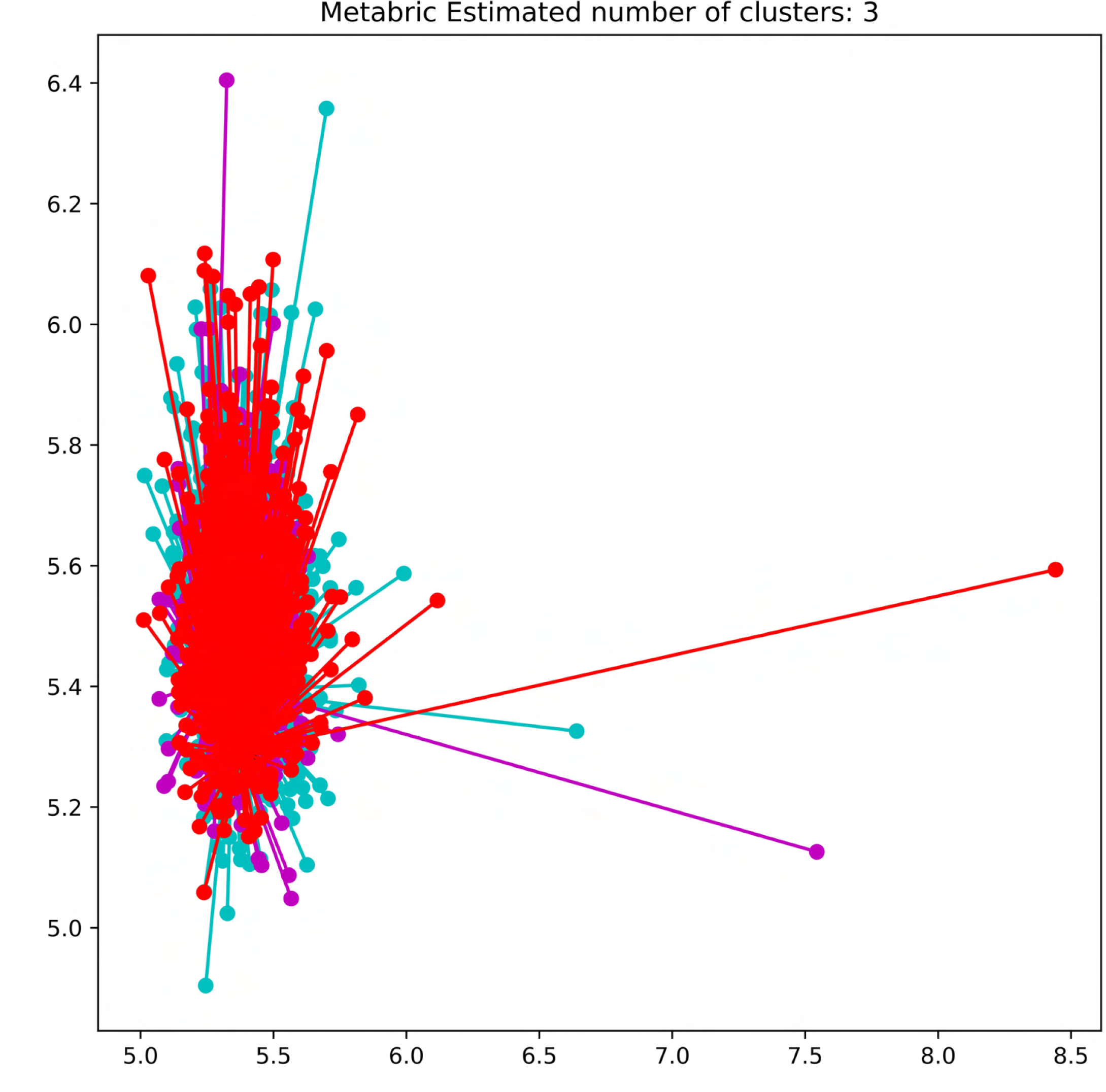

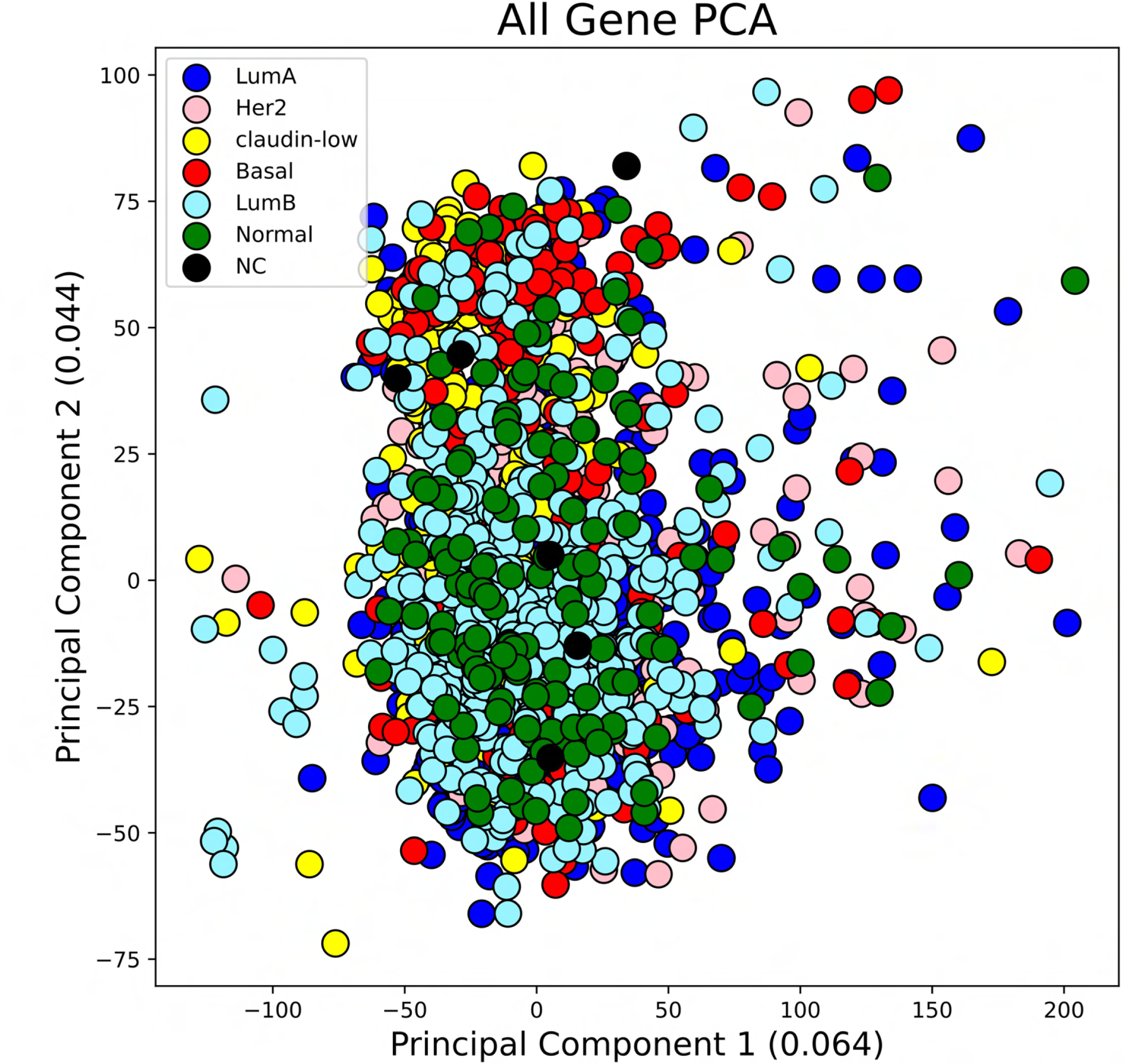

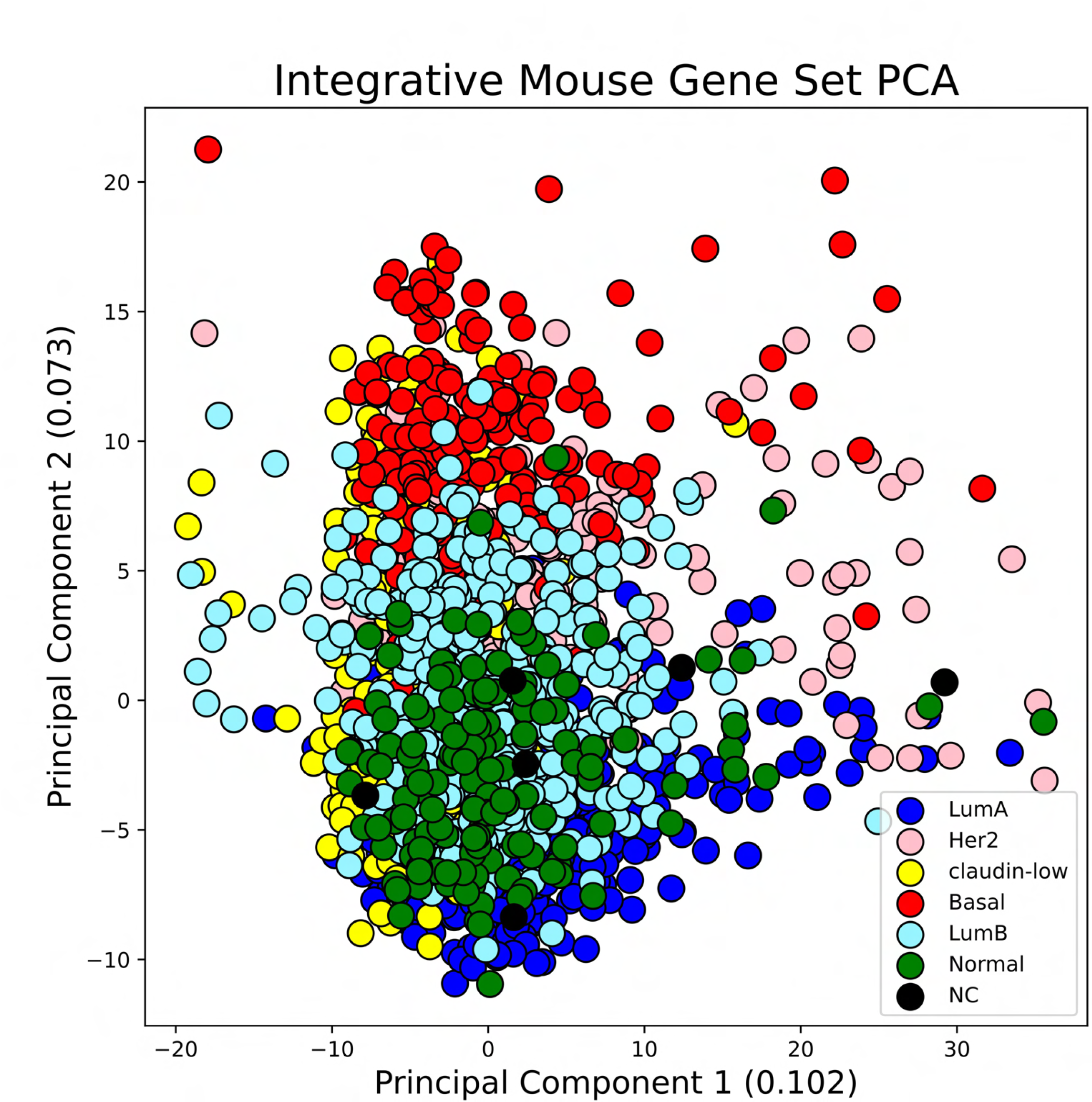

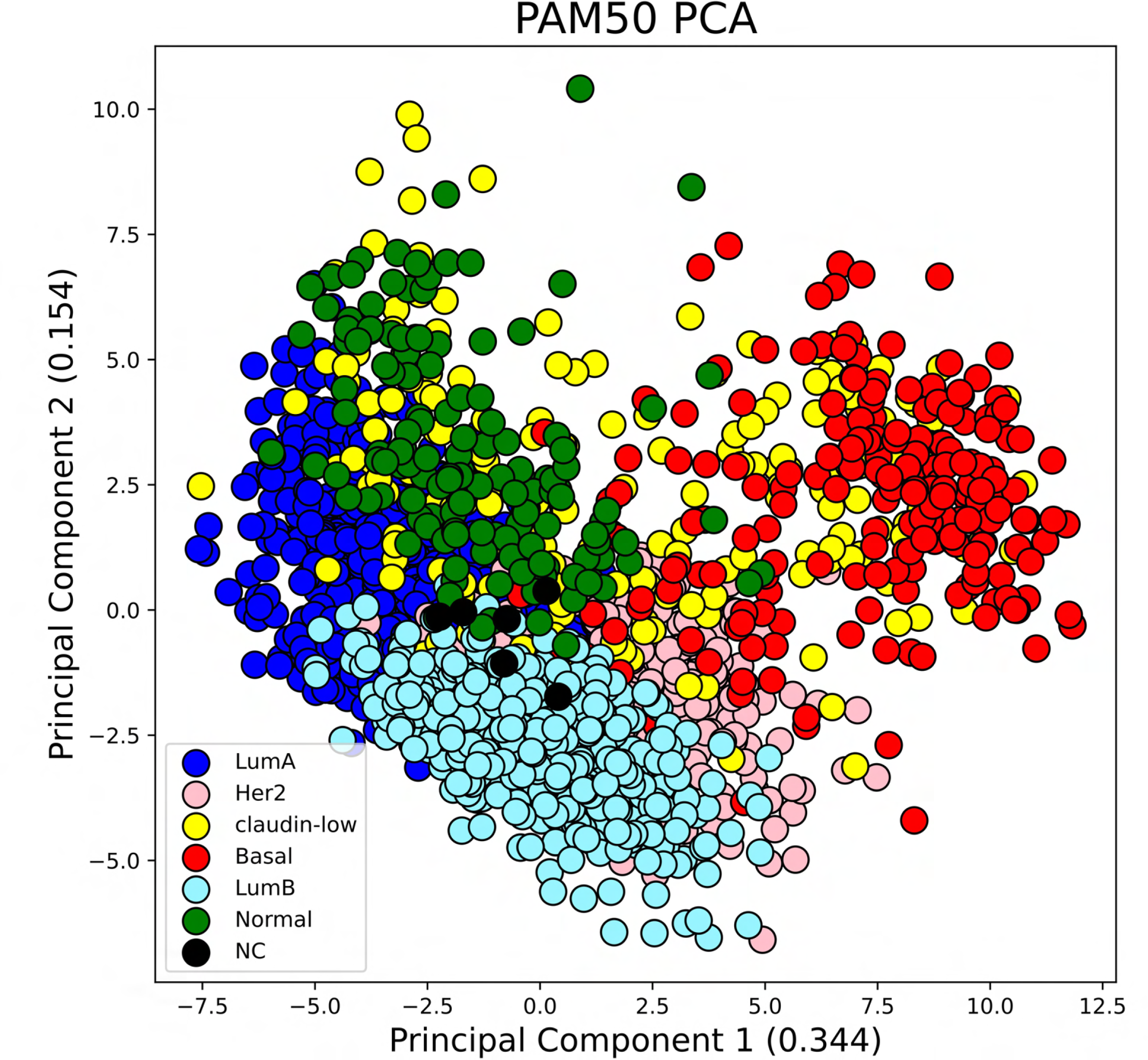

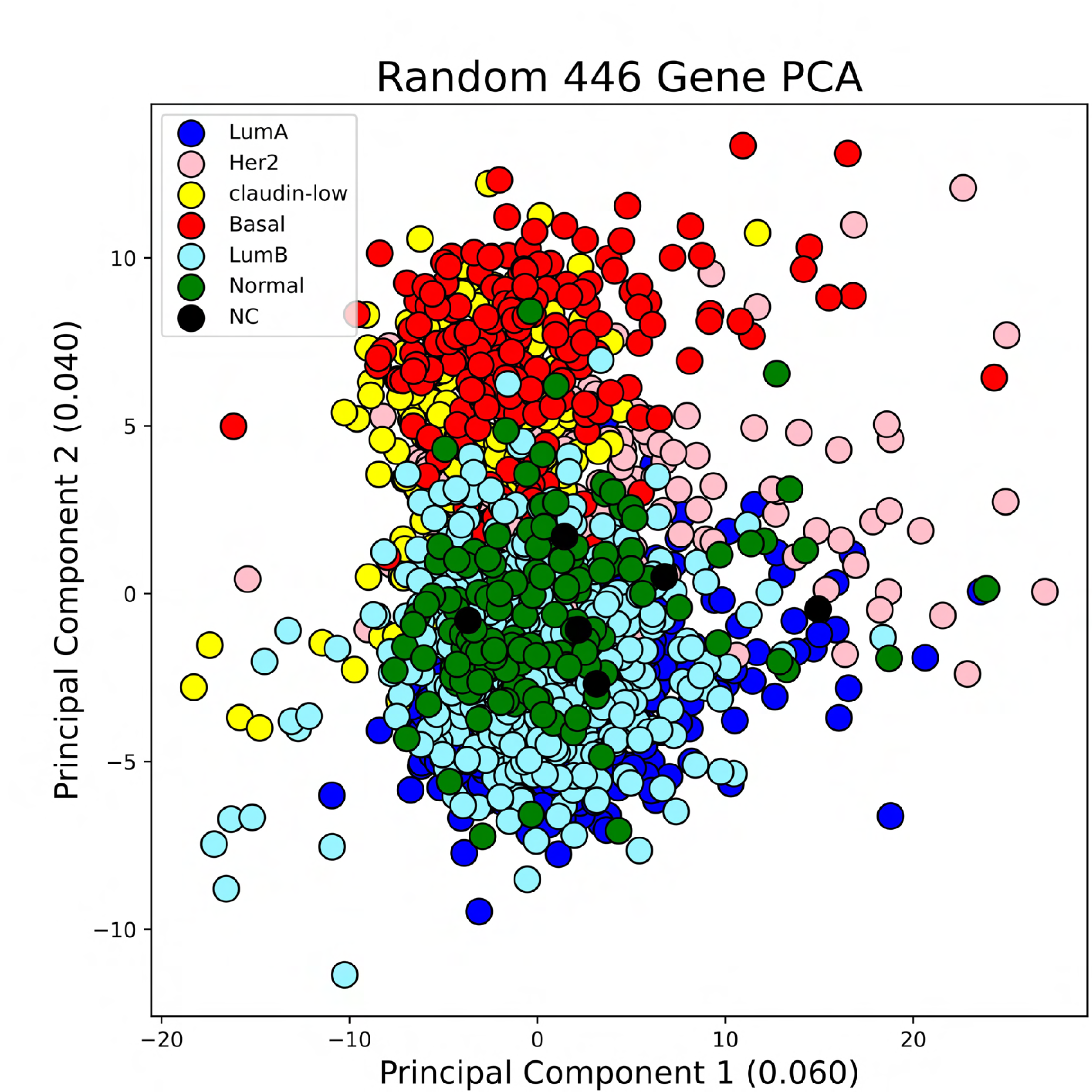

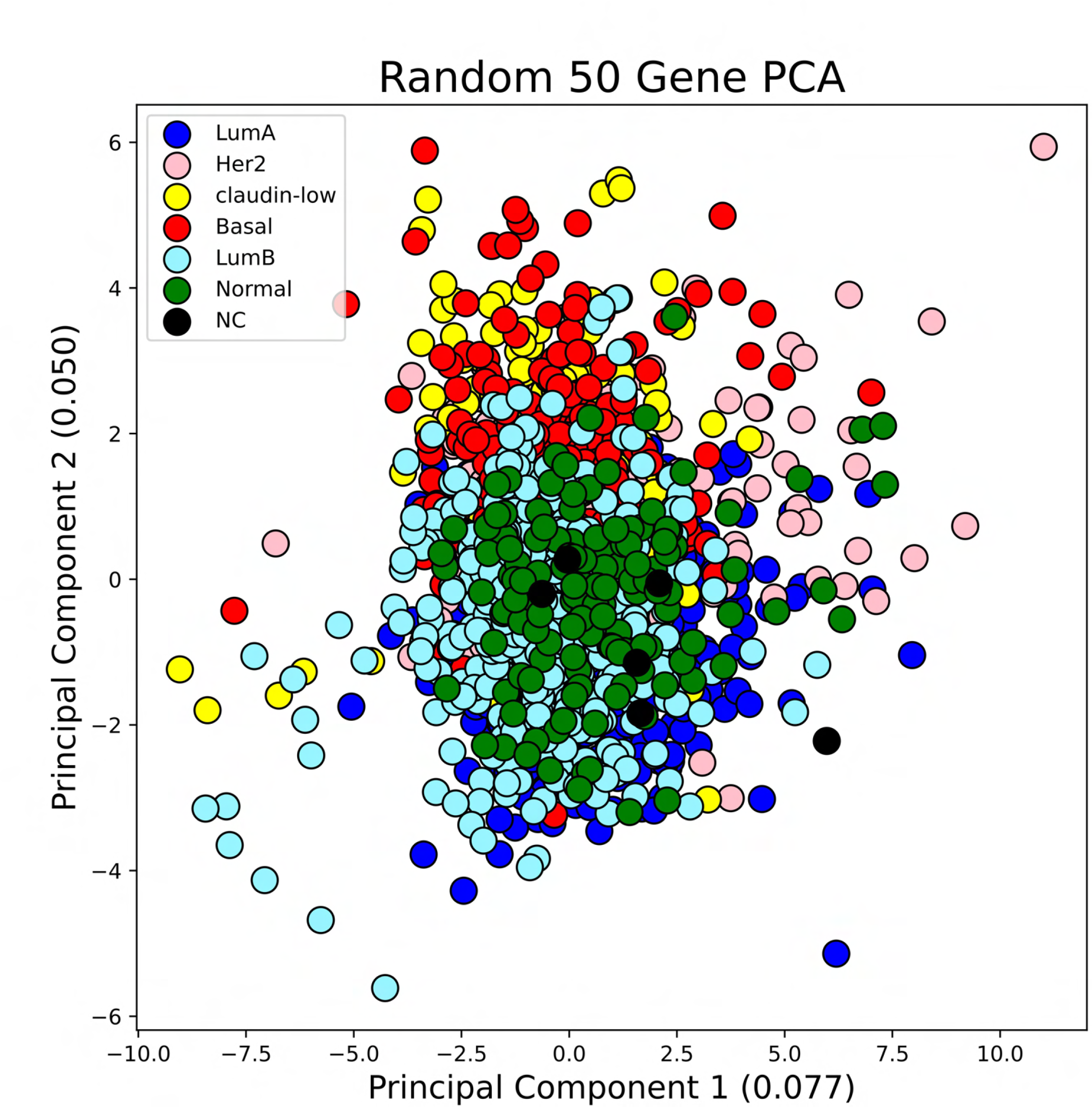

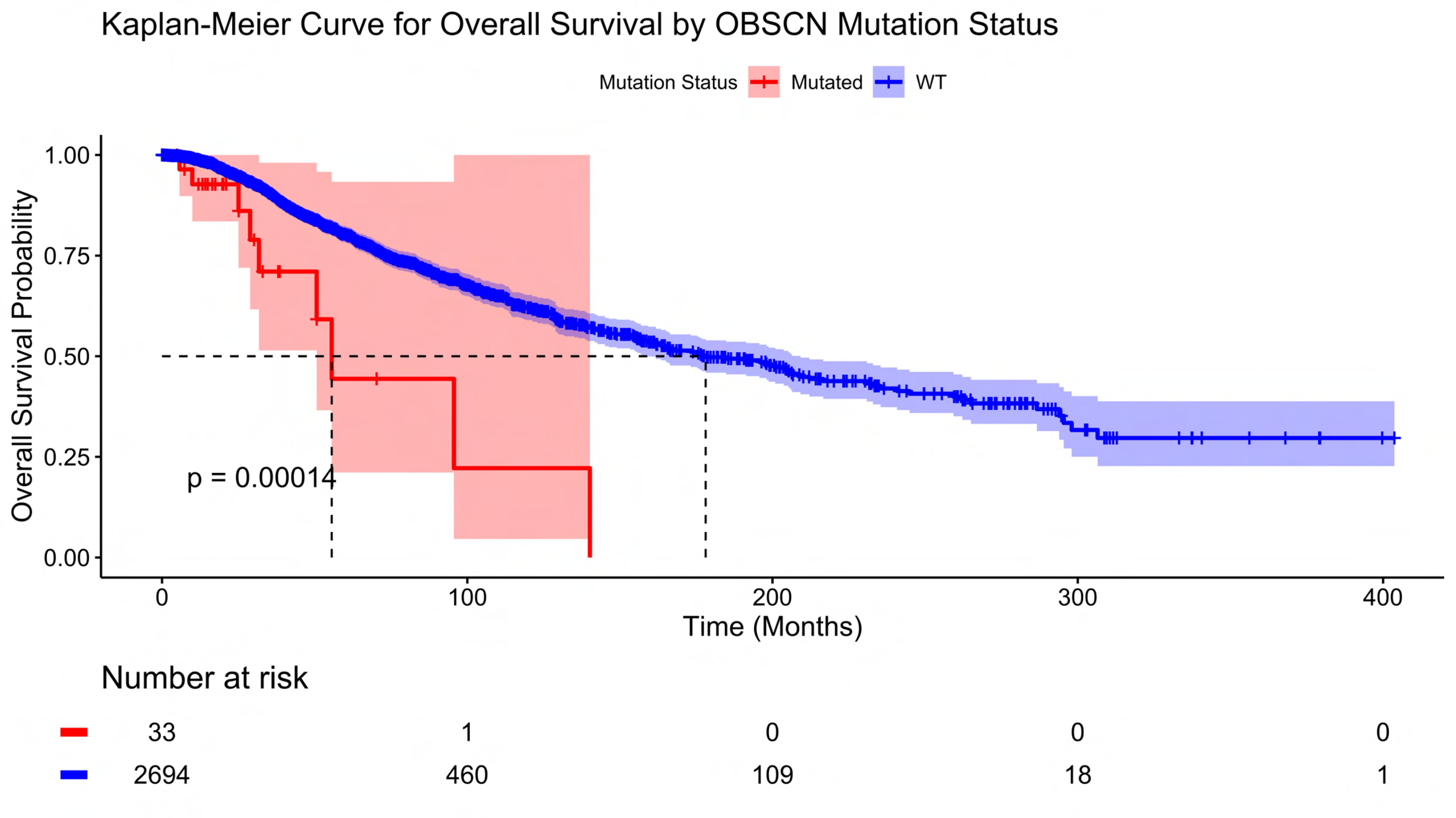

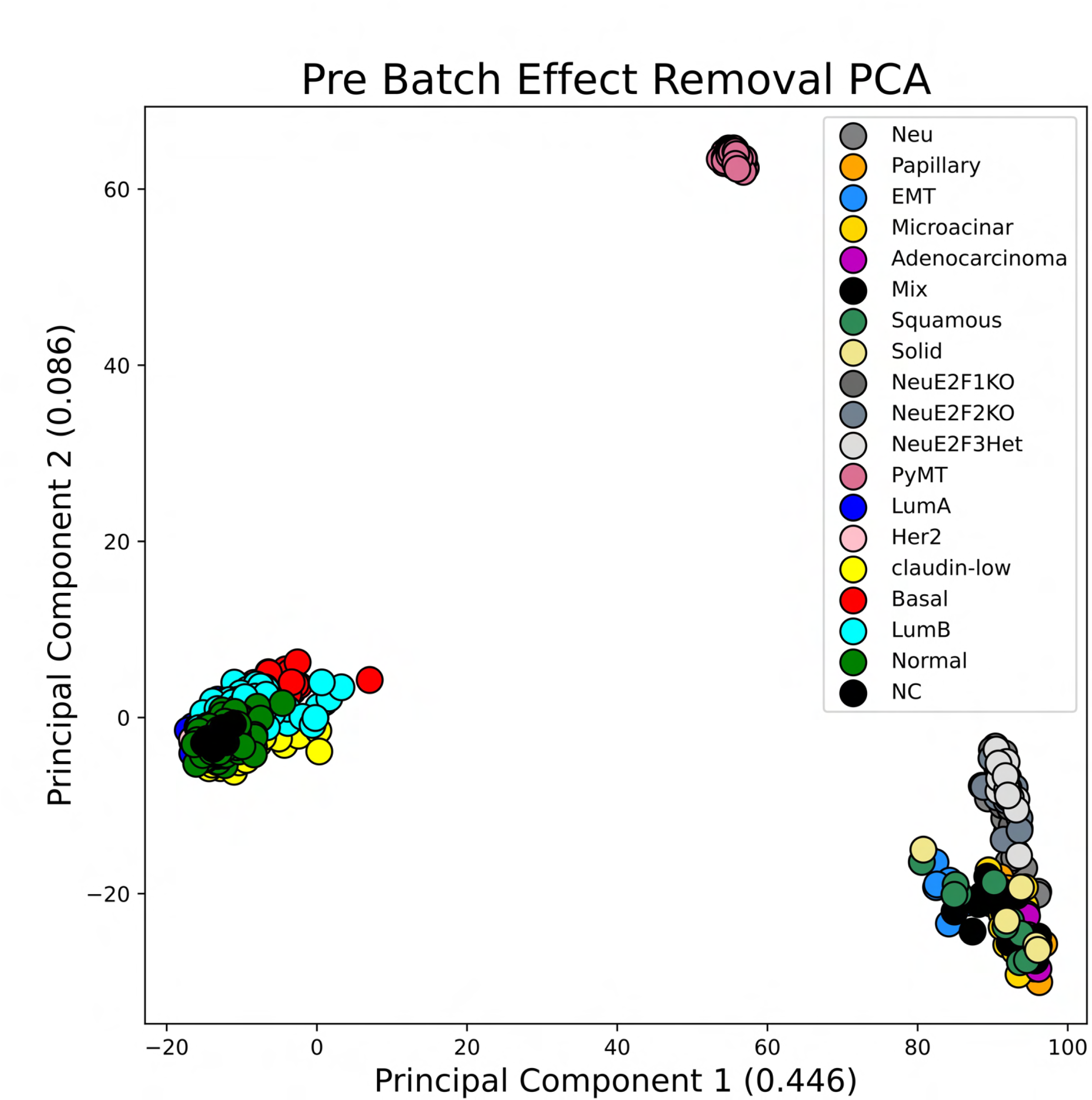

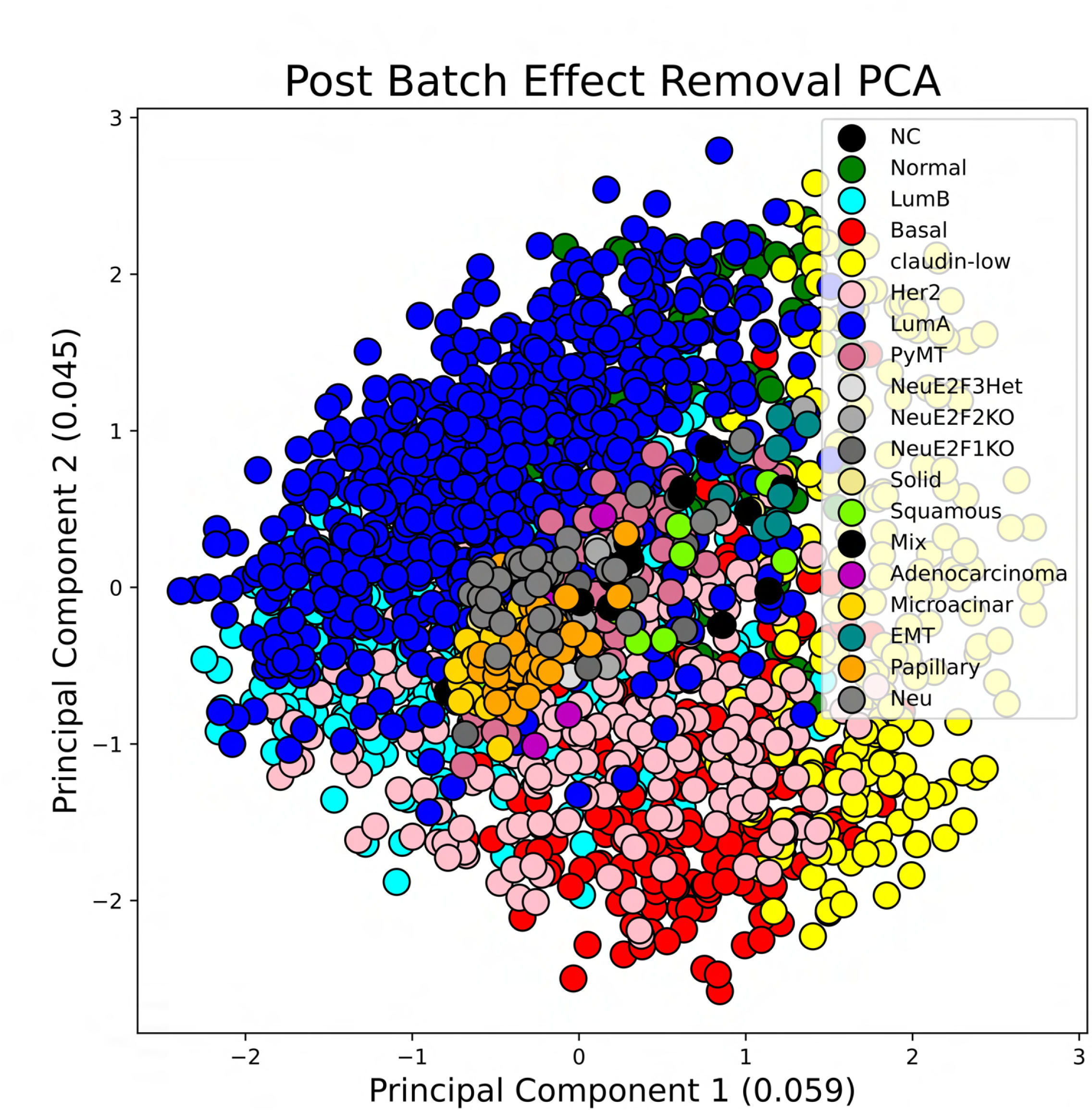

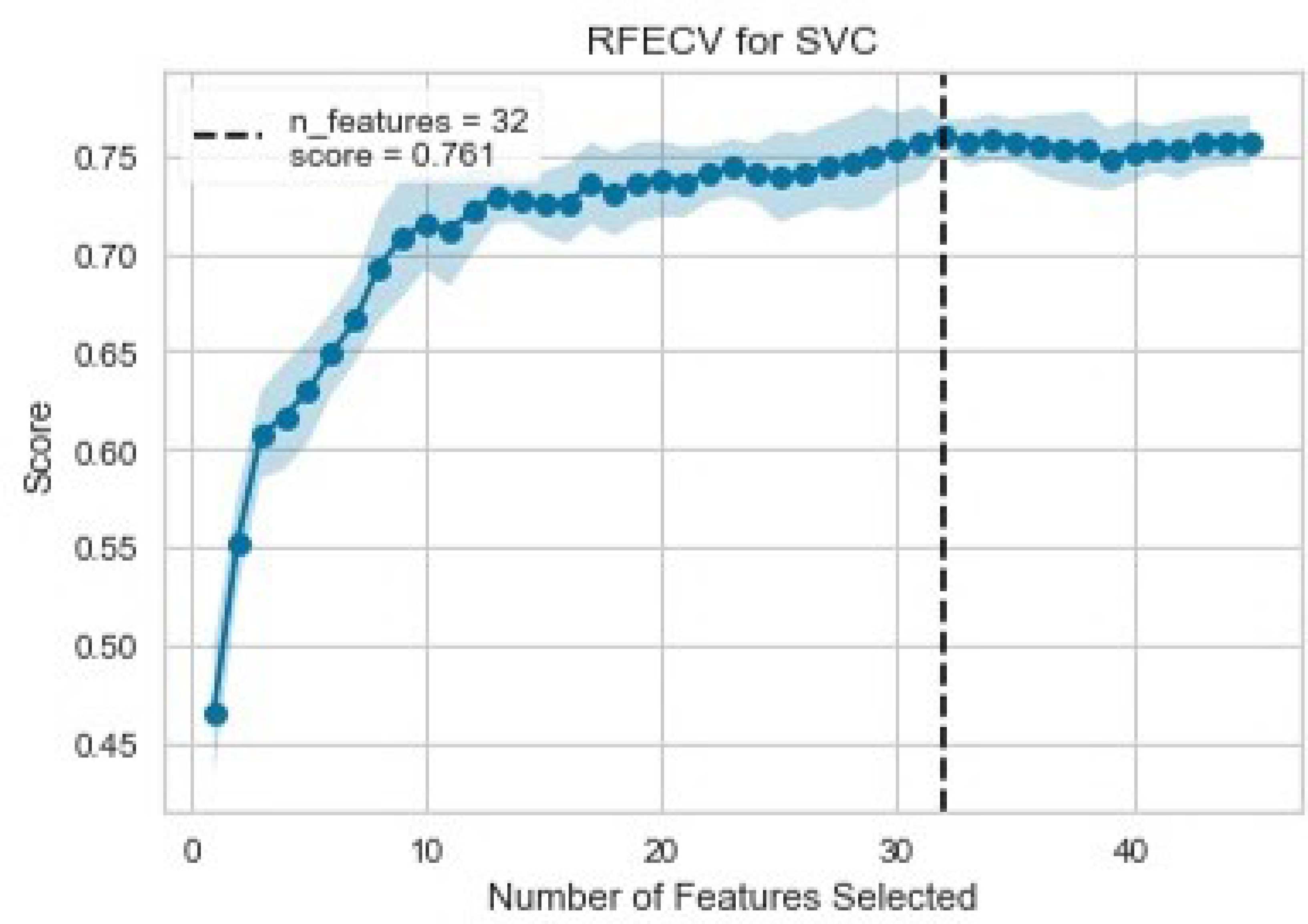

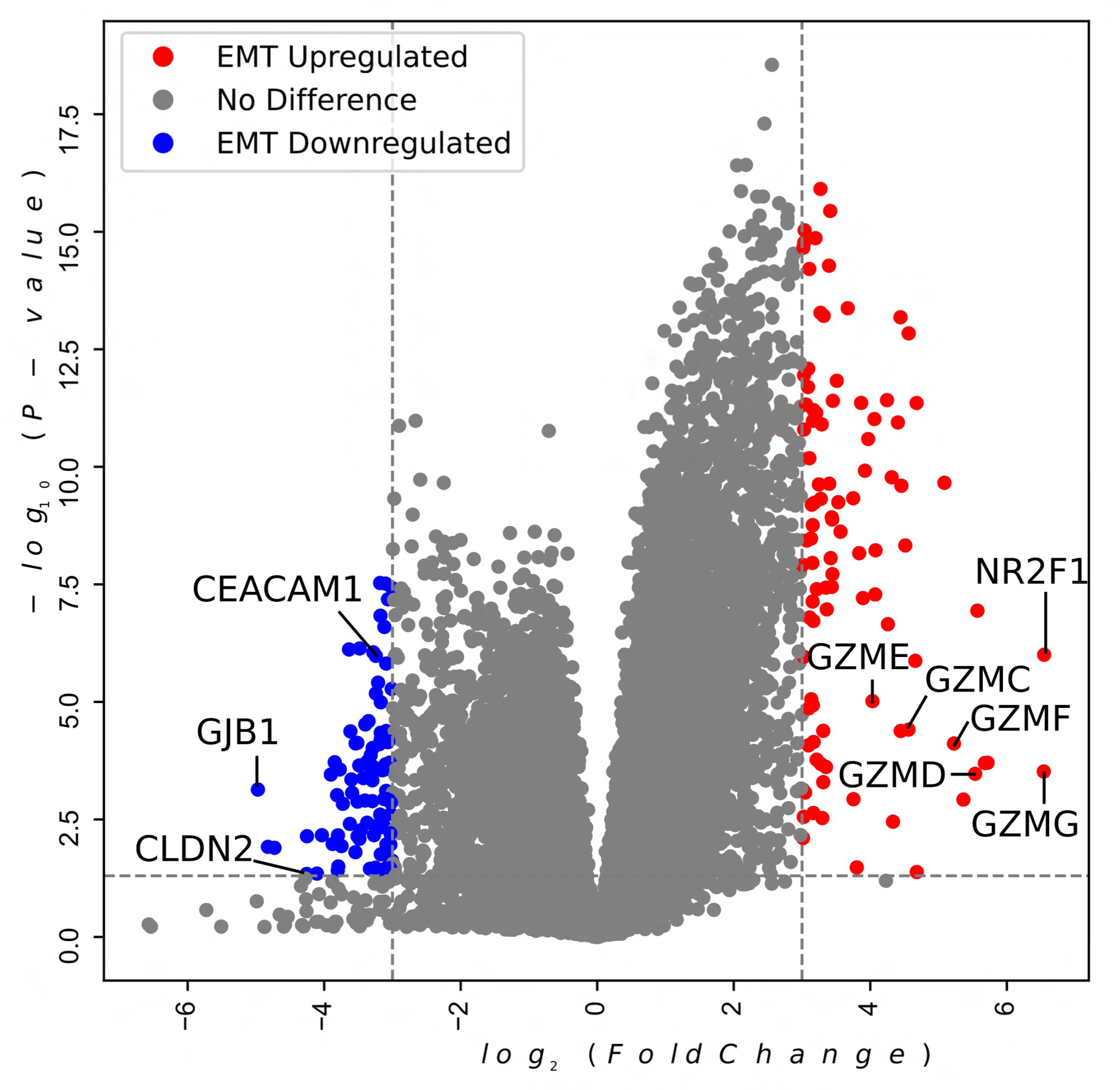

